# Short-term, long-term, and genetic determinants of human plasma proteome variability

**DOI:** 10.64898/2026.07.30.741635

**Authors:** David Benacom, Adrien Specht, Jayna C Nicholas, Robin Guillard, Madeline Gillman, Ruth Dubin, Peter Ganz, Jerome I Rotter, Kent D Taylor, Stephen S Rich, Peter Y Liu, Alexis C Wood, Michael Y Mi, Rajat Deo, Kirsi-Marja Zitting, Laura M Raffield, Charles A Czeisler, Jeanne F Duffy, Emmanuel Mignot

**Author notes:** Correspondence should be addressed to David Benacom, Adrien Specht, and Emmanuel Mignot. These authors contributed equally to this work (David Benacom, Adrien Specht).

## Abstract

Plasma proteomics is increasingly used for biomarker discovery and predictive modeling, yet diurnal protein trajectories remain insufficiently characterized. In our review of recent proteomic biomarker studies, 43% of the identified biomarkers had previously been reported to display 24-h rhythmicity. We demonstrate that ignoring these short-term dynamic effects compromises the robustness of reported models predicting health outcomes. We integrated a population-scale multi-ethnic longitudinal cohort with repeated measures over 10 years, with two cohorts of healthy adults undergoing frequent plasma sampling across days under controlled circadian, sleep and food-intake conditions. This design enabled estimation of short-term intraindividual variability (ST), long-term intraindividual variability (LT), population-level variability (POP) and genetic effects (GEN) across 7,289 protein targets. ST, LT, POP, and GEN define diverse protein trajectories, including rapid dynamics, long-term change, and individual-specific signatures. Using and generalizing this framework will facilitate covariate selection, study design, biomarker prioritization, and variability-aware modeling by users of proteomic data.

**Highlights:**

- 43% of recent reviewed plasma protein biomarkers display 24-h rhythmicity
- Short-term variability can reduce the robustness of plasma proteomic prediction models
- Short-term variability matches long-term and exceeds genetic variability
- A unified taxonomy maps 7,289 targets across four variability dimensions

**Graphical abstract:** 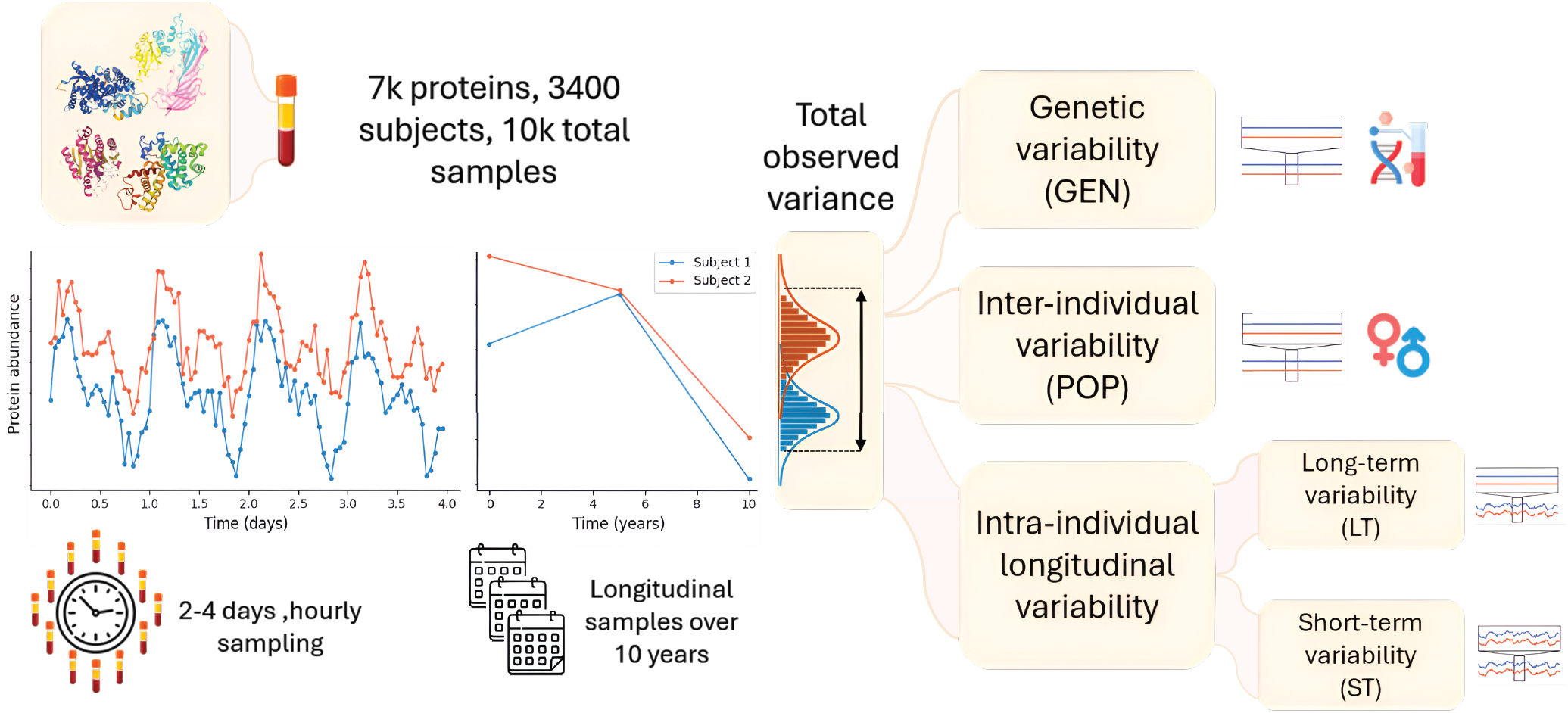

## Introduction

The identification of biomarkers is a key objective of proteomics research, with implications for diagnostics, prognostics, and the advancement of precision medicine^1,2^. In recent years SomaScan, Olink, and other multiplexed assay platforms have enabled large-scale plasma studies of general health and specific diseases^3,4^. Although these technologies have been transformative, not all probes (antibody-based for Olink, aptamer-based for SomaLogic) used for detection have established specificity, and by design, detection only interrogates a specific portion of the protein, thus can be sensitive to isoform or degradation products. This is reflected by the rather weak correlation across assays for many protein measurements^5^. SomaScan uses an aptamer-based platform with fluorescent array quantification. In its latest iteration, the assay reports on the detection of about 10,000 proteins, in three dilution classes, with assayed concentrations ranging from micro to femtomolar scale. This technical improvement in protein measurement is paralleled by advances in modeling^6^ which have enabled the development of candidate targeted panels with potential for clinical translation^7–14^. Clinical utility is however contingent on reproducibility, which must be considered in the context of both biological and technical variability^5,15^. Understanding the origin of this variability is fundamental to distinguishing disease-related differences from physiological variation or technical noise. Each technology used to measure protein abundance from blood plasma or serum has associated technical coefficients of variation (TECH) and specific technical caveats^16^. TECH can be estimated from pre-analytical and analytical procedures^17–20^ and prior studies have focused on comparing proteomics platforms for their measurement reliability^5,21,22^.

Biological variability, on the other hand, can be broadly divided into two main categories. The first is population level inter-individual variability (hereafter referred to as the population variability, POP) which includes biological sex^23^, genetics (GEN), and persistently acquired determinants, including latent infection status, early-life exposure and long-standing environmental exposures that remain stable over the sampling window. The second is intraindividual, longitudinal variability which can be further classified according to its timescale: long-term variability (LT) spanning months to years, and short-term variability (ST) occurring over hours to consecutive days. These intraindividual sources of variability include, among others, aging, seasonality, diseases, fasting-feeding, physical activity, posture, environmental factors, sleep effects, and endogenous circadian rhythms. Inter-individual variability, POP, has been associated with metabolites, protein quantitative trait loci (pQTLs), and disease status in large studies that examined ∼5,000 proteins in single blood samples from thousands of individuals^24^. However, such studies have not considered longitudinal effects, which can inflate POP estimations, and reflect the challenge and cost of obtaining serial longitudinal measurements. Nonetheless, several longitudinal studies have examined LT to assess stability of the plasma proteome across months to years, usually with two or three visits^25–27^. While most circulating proteins remain stable in adults over this timescale, LT proteins are of specific interest as they may reflect processes such as aging, chronic disease progression, medical treatment, long-term lifestyle modification, or shifts in blood cell composition^28,29^. The topic of aging has recently been approached by examining protein-based aging clocks^30–32^, which has revealed that plasma proteins do not follow a simple linear trajectory over the lifespan, thus requiring precise characterization of the plasma proteome across ages in order to characterize the effect of aging. More recently, this life-course view has been extended to early development, with longitudinal measurements from childhood through early adulthood, examining ∼5,000 proteins in individuals between the ages of 4 and 24 years, thus providing additional insight into the maturation of the plasma proteomic compartment^33,34^.

Characterizing ST is challenging because it requires longitudinal sampling at high temporal resolution over several days. ST can be modulated by physiological and behavioral factors, including fasting^35^, physical activity^36–39^, as well as procedural aspects inherent to longitudinal protocols^40–42^. While many studies routinely control for fasting as a binary covariate, reporting and standardization of sampling time remain inconsistent; failure to account for time-of-day effects is expected to reduce reproducibility and sensitivity in biomarker discovery^43,44^, and may confound disease associations when sleep or circadian physiology differs secondary to disease. To address these issues, we first examine published proteomic studies for evidence that ST-sensitive proteins are used as biomarkers and quantify the impact of ST on a published organ age prediction model. We then leverage two chronobiology datasets that provide dense repeated sampling across structured conditions, including baseline, control nights, Constant Routine, and recovery periods^45–48^, together with the Multi-Ethnic Study of Atherosclerosis large-scale population dataset^49^ (MESA), to jointly estimate ST, LT, POP and GEN in a unified framework for plasma proteome characterization.

## Results

### Protein biomarkers reported in recent large-scale studies frequently exhibit diurnal rhythmicity

We first conducted a non-systematic review to identify representative recent large-scale studies reporting clinically relevant proteomic biomarkers published in the last 5 years. From these, we identified biomarkers that have been used in prospective models for either classification of disease state, survival rate, or aging (Supplementary Table 1). We compared this list of biomarkers with the list of 1063 diurnal or circadian proteins derived from the analysis by Specht and colleagues^50^ of two circadian studies (Study 1 and Study 2, respectively S1 and S2, described in Methods) using the SomaScan 7K panel. Of 100 unique protein biomarkers identified in the non-systematic review and also present in the SomaScan 7K panel, 43 proteins were diurnally rhythmic (43%) including NT-proBNP (NPPB), Growth/differentiation factor 15 (GDF15), matrix metalloproteinase-12 (MMP12), prostaglandin-H2 D-isomerase (PTGDS), and WAP four-disulfide core domain protein 2 (WFDC2) (See Supplementary Note 1 for the full description of the studies and proteins identified in Supplementary Table 1). Several of those were already known to be diurnal, such as NT-proBNP that is associated with diurnal oscillation of blood pressure^51,52^. We then investigated whether the studies reported the time of day at which the blood samples were drawn and if that was considered in the analysis. Surprisingly, while most of the studies controlled or reported fasting, most did not report precisely the time of blood drawing, nor did they take this variable into account in their analysis. To provide an objective quantitative metric, we used a list of 217 proteins that have been approved as reliable biomarkers by the Food and Drug Administration (FDA)^53^. Among these, 166 were measured in the SomaScan-7K panel. When examined, 42 of the 166 (25.3%) were identified as having a 24-hour rhythm, including 12 (7.2%) being circadian^50^. Of note, we could not compare the effect sizes found with those obtained for diurnal fluctuations, as this must be done at the individual protein level and is not easy to extrapolate when working with complex models using multiple proteins with coefficients. Indeed, at the level of single biomarkers, the effect size of the disease might be on a different scale than the ST. For instance, in the cerebrospinal fluid (CSF), there is a 10% diurnal variation of the orexin/hypocretin immunoreactivity signal from peak to trough that is negligible compared to the many-fold differences between patients with narcolepsy and controls^54,55^. However, for complex models, a disease effect size that is smaller than ST might remain undetected. In addition, for multivariable biomarker panels, temporally structured variation across several proteins could propagate through the model and affect discrimination or calibration, particularly when sampling-time distributions differ between experimental groups.

### Short-term variability of biomarkers affects accuracy of organ aging prediction models

To demonstrate the relevance of ST in complex biomarker studies, we applied organ aging prediction models at each time point in the S1 and S2 cohorts, comprising 21 participants undergoing hourly blood sampling with plasma proteomic measurements every 2 h over multiple days in protocols originally designed to capture endogenous circadian rhythms (Table 1). The longitudinal protocol in S1 includes a baseline day, a Constant Routine (Semi-recumbent posture in dim light, continuous wake, hourly snacks across the 24-h day), and a recovery day. S2 had a Constant Routine only. The Constant Routine protocol lasts for at least 36 hours and allows assessment of protein fluctuations driven by endogenous circadian rhythms while also capturing the impact of acute sleep deprivation. Regular meals were provided on the baseline and recovery days, while 24-h caloric and fluid needs were provided as hourly snacks throughout the Constant Routine to reduce the impact of rhythmic fasting-feeding patterns on protein levels.

**Table 1.** Main characteristics of the cohorts used in this study to determine short-term (ST), long-term (LT), inter-individual population (POP), and genetic variability (GEN).

| Study | Subjects | Samples | Females | Males | Age | BMI | Version |
| --- | --- | --- | --- | --- | --- | --- | --- |
| MESA\|E1 | 5149 | 5149 | 2662 | 2477 | 62±10 | 28±5 | SomaScan 7K |
| MESA\|E4 | 4162 | 4162 | 2145 | 2017 | 66±10 | 28±6 | SomaScan 7K |
| MESA\|E5* | 3398 | 3398 | 1794 | 1604 | 69±9 | 29±6 | SomaScan 7K |
| S1<sup>#</sup> | 12 | 518 | 9 | 3 | 26±4 | 22±2 | SomaScan 7K |
| S2<sup>#</sup> | 9 | 152 | 3 | 6 | 22±2 | 23±2 | SomaScan 5K |

\*: 4332 MESA subjects with $\geq 2$ visits (mean 2.74) were used for LT. #: Note that compared to our previous study<sup>50</sup>, S1 and S2 cohorts were extended to include 4 supplemental participants. 10 MESA exam-1 participants have no recorded sex and are counted in Subjects but in neither sex column. Age and BMI are mean ± SD.

Organ age was estimated using the *organage* package^56^ (See Methods). Kidney, brain, and immune were the most stable in terms of age predictions, while heart, artery, adipose, and muscle were the most variable across the protocol duration (Fig. 1a). Predicted heart age during the 2-4 days of our protocols showed within-participant fluctuations spanning nearly two standard deviations, an effect size of the same magnitude as the threshold reported as predictive of heart disease in the study by Oh et al.^56^. In contrast, kidney age remained stable across the same time frame (Fig. 1b). Examination of the protein features underlying the heart aging model revealed that NT-proBNP, a key predictor, displayed strong short-term variation in the S1 and S2 cohorts, contributing to the inflated variability in predicted heart age (Fig. 1c). Therefore, we sought to determine whether NT-proBNP also varies with aging, which would explain its selection by the *organage* model. For that purpose, we used MESA^49^ that comprises more than 4,000 individuals of various ethnicities, ages, and disease status, with multiple longitudinal SomaScan plasma measurements available over a period of 10 years (Table 1). We found that indeed NT-proBNP abundance increases with aging, but its short-term variability is of comparable magnitude to this long-term change (Fig. 1c, third and last column). This observation suggests that proteins can exhibit distinct regulatory patterns in the short- and long-term, supporting the need for rigorous control when building predictive models with such proteins. Therefore, we sought to characterize ST, LT and POP for all proteins in the three cohorts, S1, S2, and MESA.

**Figure 1.**
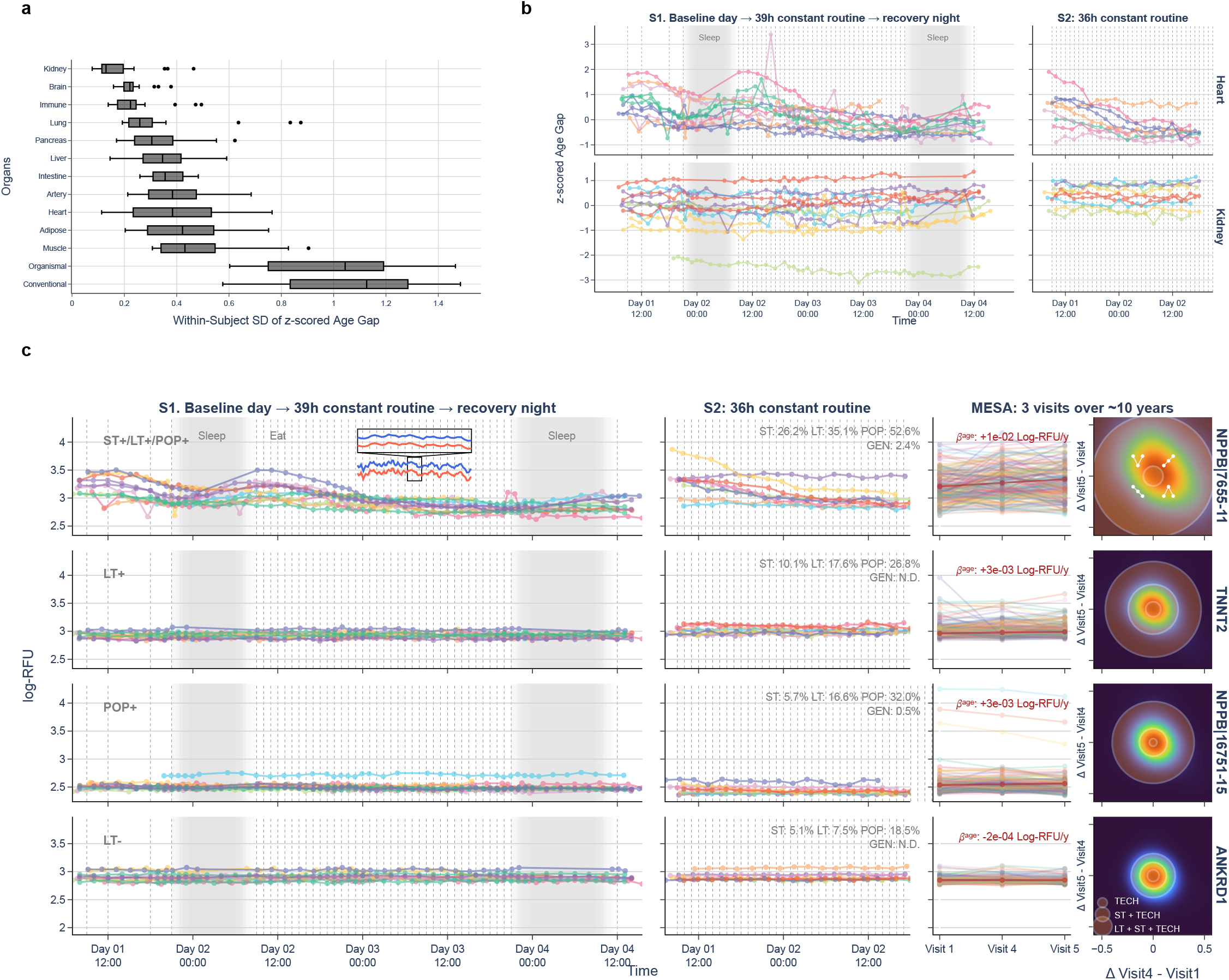
Impact of short-term variability on organ-age prediction model. Organ age was computed at each longitudinal timepoint using *organage* prediction models. **a.** Distribution of within-individual organ age-gap for cohorts S1 and S2 during the experimental protocol. Each row shows the distribution of within-individual SDs for organ-age gaps for each organ or for the aggregate of all plasma proteins (Organismal and Conventional). **b.** Participants Z-scored age gap profiles for Heart (top) and Kidney (bottom) during Constant Routine protocols. The x-axis indicates day and hour of protocol at which each sample was collected. **c.** Abundance levels of the 4 most important proteins identified by Oh et al.^56^ in the heart age model. For each protein, longitudinal plasma abundances in log relative fluorescence unit (log-RFU) are displayed for S1, S2, and MESA in the first three columns, with participants indicated by color. For S1 and S2, the x-axis indicates the day and hour of the sampling protocol; shaded regions denote scheduled sleep opportunities, and vertical dashed lines indicate meals during baseline and recovery days and snacks during the Constant Routine protocol. S1 (n = 12; 518 samples) was collected across a baseline day, a 39-h Constant Routine protocol, and a recovery night, whereas S2 (n = 9; 152 samples) was collected during a 36-h Constant Routine protocol. For MESA (n = 3,398; three visits per participant over 10 years), the x-axis indicates visit number; 200 participants were subsampled for plotting purposes only. The final column shows the 2D distribution of individual between-visit differences in MESA, providing a quadrant-based classification of longitudinal patterns and a visual decomposition of observed long-term variability into ST, LT, and TECH components.

### Estimation of short-term, long-term, population, and genetic variability, across 7,289 SomaScan protein targets in the human plasma

To estimate ST, LT and POP across the 7,289 SomaScan protein targets, we fitted one linear mixed model per aptamer across the three cohorts. The model partitioned variability into short-term residual fluctuation, long-term visit-to-visit drift, and inter-individual differences after accounting for TECH and fixed covariates. For genetic variability (GEN), we computed the cumulative variance explained by sentinel pQTLs found using genetic data from the 5,149 included MESA participants with protein measures available at exam 1 (See Methods for additional details). 3,992 probes had at least one significant pQTL: 1,456 cis only (within 1 Mb of transcription start site; p < 5 × 10⁻^8^), 1,671 trans only (p < 6.8 × 10⁻^15^), and 865 with both (Supplementary Figs. 1-4). Polygenic scores (PGS) were derived in MESA, using the effect sizes from sentinel pQTL as weights. Each PGS was regressed on the observed protein abundance; the resulting R^2^ was used to estimate a GEN value, representing the percentage of total variance explained by genetics. These values were highly correlated with previously reported SomaScan genetic variance estimates (Supplementary Fig. 5). Of note, for probes without any pQTL detected, GEN was reported as 0 for mean/median comparison, and as non-determined (N.D.) in the subsequent figures, although it is possible that these proteins have genetic contributions to circulating abundance that could have been detected with a larger cohort size. The full list of proteins and corresponding estimates is available in Supplementary Table 2, with detailed genetics analysis in Supplementary Table 3.

Mean values of the estimated variability components were 11.0% for ST (Median = 7.5%; upper quartile, ST+ > 13.5%), 14.8% for LT (Median = 11.3%; upper quartile, LT+ > 16.9%), and 27.5% for POP (Median = 23.7%; upper quartile, POP+ > 30.5%). PGS for each probe explained on average 4.0% of the variance in circulating probe measures (GEN; Median = 0.3%; upper quartile > 3.6%). Given the skewness of GEN values, GEN+ was defined as probes for which total variance explained by genetics was > 20%. The total number of aptamers in each category and their overlap are represented in Fig. 2a. Positive correlations were observed between LT/POP, POP+/GEN+ and ST+/LT+ (Fig. 2b,c); some of these associations were non-linear, with LT and ST correlating primarily in their upper ranges, whereas POP and LT correlated at their lower ranges (Fig. 2c).GEN+ probes were highly correlated with POP+ (Fig. 2c,d). TECH was strongly correlated among SomaScan assay versions, but only weakly with ST, LT or POP (|ρ| ≤ 0.22), confirming that TECH is more likely to represent aptamer-binding characteristics.

**Figure 2.**
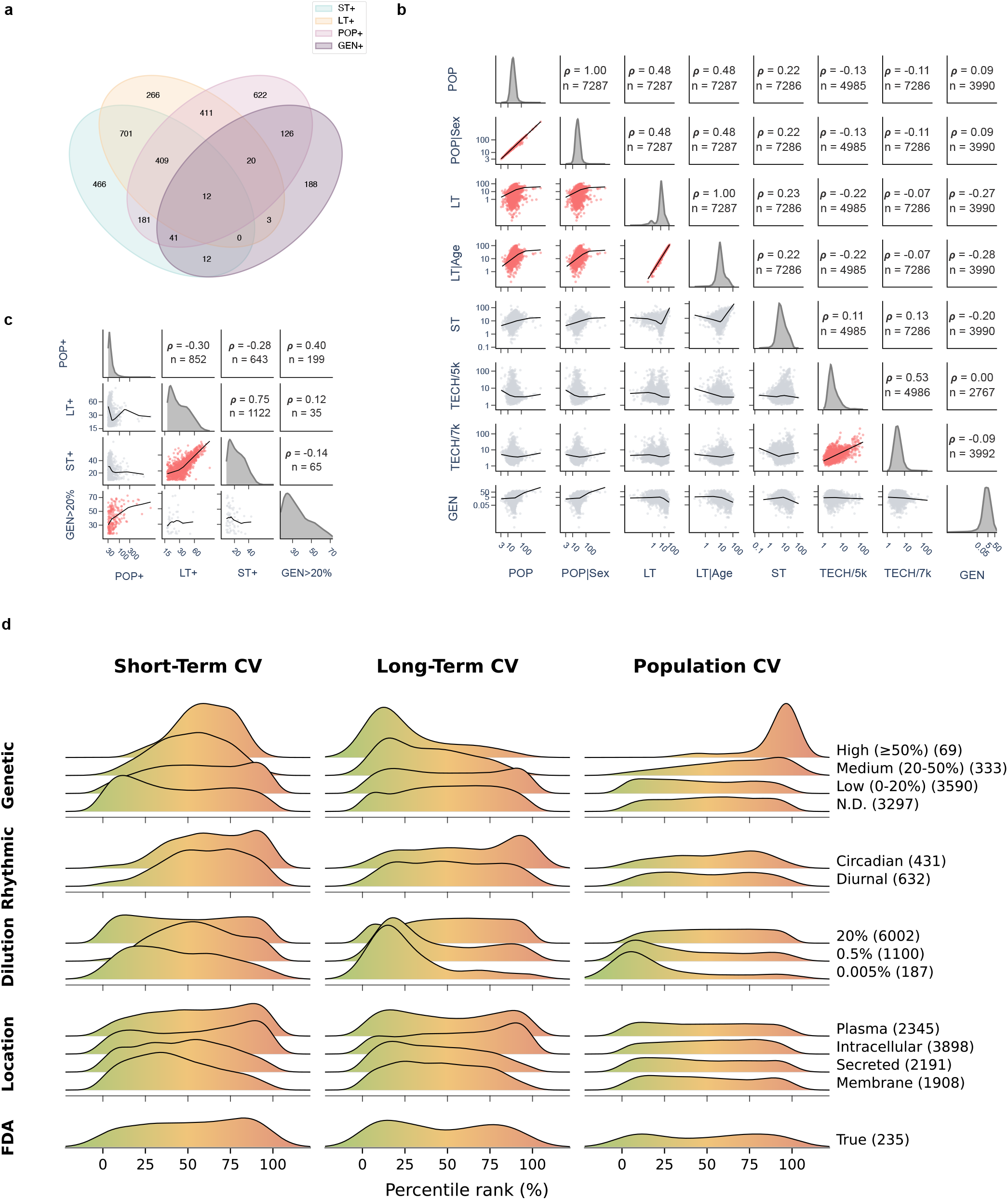
Characterization of the ST, LT, POP and GEN metrics. **a.** Venn diagram depicting the intersection of ST+, LT+, POP+ and GEN+. **b.** Pairwise relationship between stability metrics. Upper triangles report Spearman correlation coefficients (ρ) and n, number of proteins. Lower triangles show bivariate scatterplots colored by the sign of the correlation (red=positive, blue=negative), with LOWESS curves in black. Diagonal panels display the marginal distributions for each metric. Metrics include population CV (POP), Sex-adjusted POP (POP|Sex), long-term CV (LT), short-term CV (ST), intra-plate technical CV estimated by SomaScan for their assays (TECH/4 or /4.1) and GEN, representing variability explained by pQTLs. **c.** Filtered on highly variable proteins, i.e. > Q75. **d.** Ridgeline plots show the percentile-ranked variability of 7,289 SomaScan protein targets for ST, LT, and POP. Rows correspond to five annotation groups: computed GEN category, rhythmicity^50^, dilution class provided by SomaScan, cellular localization from the Human Protein Atlas^57^ (HPA) and FDA approved biomarkers^53^ (for protein, not necessarily SomaScan aptamer). Background color bands provide interpretive ranges for each metric (green: low variability, yellow: moderate, red: high).

We next asked whether the four variability components recapitulate known biological and technical features of the plasma proteome. Biologically, proteins previously identified as diurnal or circadian^50^ exhibited elevated ST, as expected, together with a bimodal distribution of LT and POP (Fig. 2d). Intracellular proteins showed positively skewed ST and LT distribution in comparison to membrane-bound or secreted proteins, while subcellular localization did not meaningfully associate with POP (Fig. 2d). When stratified by SomaScan dilution class, from 0.005% (highly abundant) to 20% (extremely low abundance), highly abundant proteins displayed lower ST, LT, and POP; this gradient weakened in the intermediate 0.5% class and disappeared at 20% (Fig. 2d).

Proteins previously associated with aging waves^31^ showed ST comparable to the panel as a whole, but reduced LT and POP (Extended Data Fig. 1). These protein associations were derived largely from the INTERVAL cohort^58,59^, sampled from blood donors without fasting or time-of-day standardization, implying that short-term variability cannot be accounted for without metadata. This indicates that these markers could carry a short-term component that single-sample designs cannot capture. This is worth accounting for as systematic bias could be associated with age of participants and time of blood draws for instance. We also examined selected predictors of organ-specific age^56^ for different organs (Extended Data Fig. 1). However, these models are weighted combinations of proteins, so their feature sets cannot be interpreted as flat categories. The heart model illustrates this with ST values of its features that are low overall, yet its most heavily weighted predictor, NT-proBNP, was the highest-ST protein in the set (ST = 26.2%) leading to the variability in organ age identified previously (Fig. 1b).

We further evaluated whether the choice of proteomics platform, or the performance of the platform, influences the types of variability that can be captured. Probes with high SomaScan–Olink correlation (suggesting the ability of both platforms to consistently measure the same protein^22,53^) tended to display higher ST and POP, whereas LT showed a bimodal distribution; proteins represented in the Mass Spectrometry (MS) - Internal Standard Targeted panel from Kirsher et al.^53^, a targeted reference proteomics dataset, were generally low in both LT and POP (Extended Data Fig. 1). Finally, although we expected FDA-approved biomarkers curated by Kirsher et al.^53^ to be enriched among LT+ or POP+ positive proteins, they instead showed a slight shift toward higher ST values, without comparable enrichment in LT or POP. These results suggest that platform performance and assay design may shape which subsets of proteins are most readily captured within each variability regime, although ST, LT, POP, and GEN were estimated here from SomaScan data rather than compared directly across platforms.

### A variability-based taxonomy of human plasma proteins

To gain an intuition of how ST, LT, POP and GEN can define plasma proteins, we examined the joint distribution of ST and POP (Fig. 3a). This allows identification of proteins that can be highly dynamic within individuals during the day yet with a baseline of abundance conserved across individuals (ST+/POP−; upper-left quadrant). Conversely, one can also identify proteins that display baseline dispersion between individuals while being stable on the short-term (POP+/ST−). Surprisingly, we observed that glyceraldehyde-3-phosphate dehydrogenase (GAPDH), a housekeeping protein frequently used as an experimental control, was in the POP+/ST+ quadrant, suggesting that it varies both between individuals and at the short-term scale. With additional refinement provided by LT and GEN, we defined several categories of proteins (Fig. 3b) for which we provided examples of individual protein trajectories to illustrate how each variability profile can inform biomarker selection and study design (Fig. 4).

**Figure 3.**
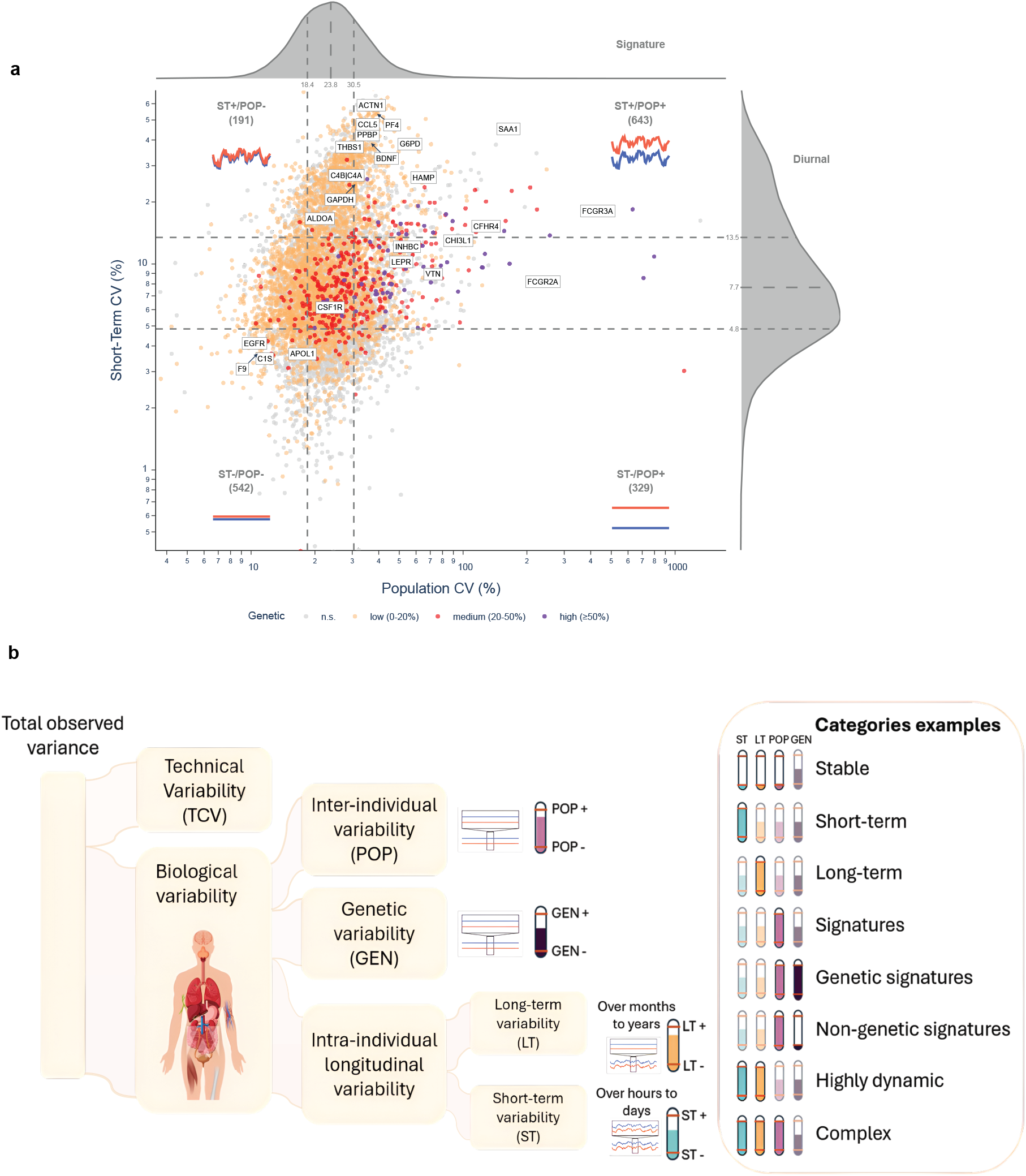
Derivation of a variability-based taxonomy of plasma proteins. **a.** Joint distribution of short-term (ST) and population (POP) variability across plasma proteins to visualize categories. Each point represents a SomaScan target, colored by the fraction of variance explained by genetic factors (unknown, low, medium, high). As expected, GEN is visually enriched among proteins with elevated POP, consistent with the contribution of pQTLs to baseline interindividual variation. The x-axis shows population-level variability (POP, %), and the y-axis shows short-term biological variability (ST %). Horizontal and vertical dashed lines mark distribution-based quartile thresholds used to define four variability quadrants: ST+/POP−, ST+/POP+, ST−/POP−, and ST−/POP+, with the number of proteins in each group shown. Kernel density plots along the top and right margins summarize the distributions of POP and ST, respectively. **b.** Schematic representation of the sources of variability measured. The technical variability (TECH) is removed from the model using quality control variables. The remaining biological variability is split between inter-individual variability (POP), including genetics (GEN) and intra-individual variability, which is further split into short-term (ST) and long-term (LT). Schematic colored gauges represent the variability in each of the categories. Based on these characteristics, we defined as examples: 1. (ST–/LT–/POP–) *Stable markers.* 2. (ST+) *Short-term marker*s. 3. (LT+) *Long-term markers*. 4. (POP+) *Signatures*. 5. (POP+/GEN+) *Genetic signatures*. 6. (POP+/GEN-) *Non-genetic signatures*. 7. (ST+/LT+) *Highly dynamic markers.* 8. (ST+/LT+/POP+) *Complex markers*.

**Figure 4.**
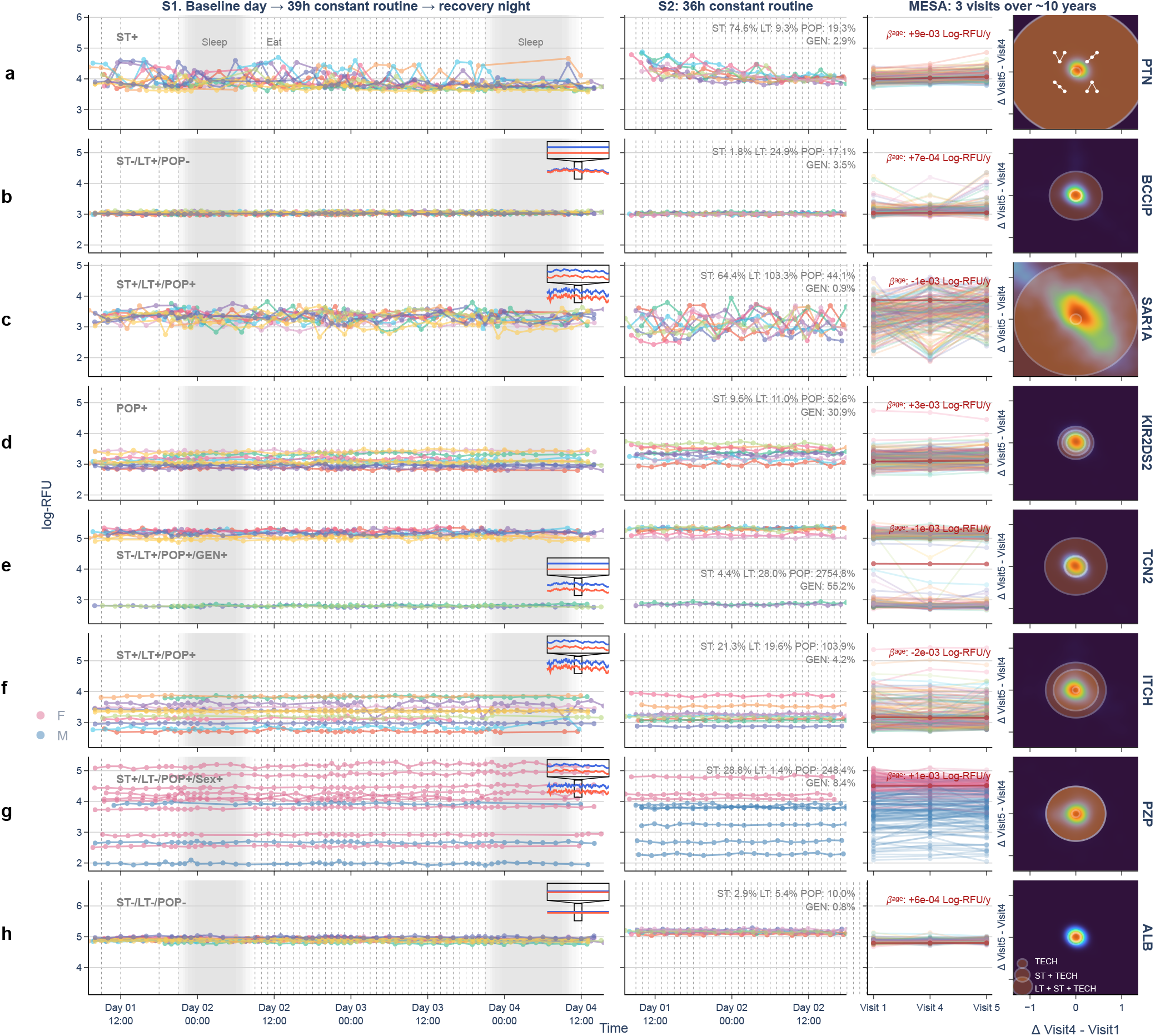
Representative plasma protein trajectories from the variability-based taxonomy. Representative protein trajectories illustrating variability profiles across short-term (ST), long-term (LT), and population-level (POP) variability. From top to bottom, proteins shown were selected for illustrative purposes: **a.** PTN archetype of *ST+* protein (here ST+); fluctuates over the short term but not over the long term or between individuals. **b.** BCCIP archetype of *LT+ specific* protein (here ST– /LT+/POP–); fluctuates long-term but neither over the short-term nor between individuals. **c.** SAR1A, ST+/LT+/POP+, *highly dynamic* protein, responsive across short- and long-term timescales. **d.** KIR2DS2, archetype of *signature* protein (here POP+/GEN+); fluctuates markedly between individuals, with a strong genetic contribution. **e.** TCN2, LT+/POP+/GEN+, fluctuates long-term and between individuals with a strong genetic influence. **f.** ITCH, ST+/LT+/POP+; fluctuates short- and long-term and between individuals despite low genetic influence. **g.** PZP, ST+/LT–/POP+/Sex+; fluctuates short-term and between individuals with a strong sex influence. **h.** ALB, ST–/LT–/POP–, *stable* protein across all timescales. For each protein, longitudinal plasma abundances (log-RFU) are displayed for S1, S2, and MESA in the first three columns, with participants indicated by color. For panel g, colors instead indicate sex. For S1 and S2, the x-axis indicates the day and hour of the sampling protocol; shaded regions denote scheduled sleep opportunities, and vertical dashed lines indicate meals during baseline and recovery days and snacks during the Constant Routine protocol. S1 (n = 12; 518 samples) was collected across a baseline day, a 39-h Constant Routine protocol, and a recovery night, whereas S2 (n = 9; 152 samples) was collected during a 36-h Constant Routine protocol. For MESA (n = 3,398; three visits per participant over 10 years), the x-axis indicates visit number; 200 participants were subsampled for plotting purposes only. The final column shows the 2D distribution of individual between-visit differences in MESA, providing a quadrant-based classification of longitudinal patterns and a visual decomposition of observed long-term variability into ST, LT, and TECH components.

As mentioned previously, *ST+* proteins may be useful for studies of circadian rhythms or short-term metabolic regulation. As such, circadian or rhythmic proteins were frequently captured as ST+ (Fig. 2d). Examples include NT-proBNP (discussed above), rhythmically secreted hormones such as thyroid-stimulating hormone (TSH) and pro-opiomelanocortin (POMC)-derived corticotrophin- lipotrophin, and meal-timing-sensitive proteins such as insulin-like growth factor-binding protein 1 (IGFBP1) (Supplementary Table 2). However, *ST+* does not necessarily imply rhythmicity. Pleiotrophin (PTN), a secreted heparin-binding growth factor involved in neurotrophic signaling, angiogenesis and tissue remodeling^60^ displayed marked short-term fluctuations (Fig. 4a), but no rhythmic effect was detected in our previous analysis^50^. Its variability is therefore unlikely to be fully addressed by sampling-time control alone.

On longer time scales, *LT+* proteins may be particularly relevant for monitoring chronic trajectories rather than for single-timepoint diagnosis. For example, olfactomedin-4 (OLFM4) displayed pronounced long-term variability in our analysis (LT = 64.6%), with limited variability between individuals and short-term variability only marginally above our ST+ threshold. OLFM4 is a measurable secreted plasma protein linked to neutrophil-mediated inflammation^61^, and reported as a potential biomarker in gastrointestinal cancer and infectious disease severity^62–64^. Its variability profile therefore suggests that it may be a promising candidate for longitudinal biomarker studies. The *LT+* category also included intracellular proteins, as noted above (Fig. 2d). For example, BRCA2/CDKN1A- interacting protein (BCCIP), a cell-cycle and DNA-damage-response protein implicated in cancer biology^65–67^, displayed long-term variability without substantial associated POP or ST (LT = 24.9%, Fig. 4b; Supplementary Table 2). These two proteins are worth examining together, as they do not share the same relationship with short-term variability and age: OLFM4 shows some short-term variation but no detectable age trend, whereas BCCIP is stable in the short term and increases systematically with age. The plasma interpretation of BCCIP remains cautious because BCCIP is not a classical secreted protein and may instead reflect cell-derived material or tissue turnover.

However, these two categories are not exclusive. *Highly dynamic* (ST+/LT+) proteins such as GTP-binding protein SAR1a (SAR1A) can display joint short- and long-term variability. SAR1A is a component of the coat protein complex II (COPII)^68,69^, which regulates intracellular shuttle of proteins between the Golgi apparatus and the endoplasmic reticulum, a process likely responsive to many metabolic signals (Fig. 4c). Such proteins require careful longitudinal design and covariate control to interpret as biomarkers or could be excluded from panels to gain statistical power.

In addition to temporal variability, *signature* proteins (POP+) can characterize individuals from a single measurement by capturing stable differences in baseline abundance and may therefore be useful for risk stratification or disease susceptibility. A classic example is provided by genetic signature proteins (POP+/GEN+), including polymorphic immune targets such as killer cell immunoglobulin-like receptor 2DS2 (KIR2DS2), which has been associated with increased risk of several immune disorders, notably HLA class I-associated diseases^70–73^ (Fig. 4d). KIR2DS2 is encoded by a gene with known copy number variation in the genome^74^, which partially explains such GEN effects, but the KIR assay signal remains complex to interpret (Supplementary Fig. 6). Another example is the cobalamin (vitamin B12) transporter Transcobalamin 2 (TCN2), which showed extreme population-level variability with GEN accounting for 55.2% of its total variability (Fig. 4e). In our dataset, a single intronic TCN2 variant accounted for most of the genetically explained variance in its baseline abundance (Supplementary Fig. 7). This strong genetic contribution is consistent with previous reports linking *TCN2* variants to circulating holotranscobalamin and cobalamin-related neurological phenotypes, as well as plasma–CSF protein balance^75,76^. Importantly, TCN2 also displayed LT+ variation that may remain biologically informative when interpreted relative to an individual’s own baseline.

Not all POP+ signatures were primarily genetic, however. For instance, Itchy E3 Ubiquitin Protein Ligase (ITCH), an endosomal protein involved in the pathogenesis of severe autoimmune disorders^77,78^, was identified as *non-genetic signature* (POP+/GEN-) with participant-specific baselines that were not fully explained by known genetics (GEN = 4.2% vs 55.2% for TCN2; Fig. 4f). Such *non-genetic signature* effects could be explained by other defining factors such as sex, leading us to also compute POP conditioned by sex. In the case of ITCH, POP|sex = 103.9% ≈ POP, suggesting that for this protein the baseline effect is independent of sex, and that could be explained by more complex factors such as persistent epigenetic factors or environmental exposures. Even for classical sexually dimorphic proteins such as Pregnancy zone protein (PZP; Fig. 4g), POP = 248.4% ≠ POP|sex = 142.5%, sex does not fully explain POP, illustrating that *signature* effects can persist in addition to the sex-related variability that is usually controlled in models.

Finally, *stable* analytes show minimal variation across individuals and timepoints and offer little biological contrast but could be useful as normalization anchors across batches. This category includes abundant, homeostatic proteins such as those in the complement and coagulation cascades. However, we note that apparent stability can be a measurement artifact, since proteins whose true variation is small relative to the assay’s detection and saturation limits retain little signal once TECH is removed and appear artificially stable. This affects both ends of the dynamic range with near-saturation high-abundance aptamers such as albumin (ALB; Fig. 4h), and near-detection-limit probes so the category is enriched for hard-to-measure proteins as much as truly invariant ones, and the two are best distinguished by signal quality rather than the variability estimate alone.

Indeed, not all structured variability should be interpreted as intrinsic biological regulation without further control. While we accounted for technical variability related to SomaScan measurements, we also identified protocol-related and pre-analytical sources of variation that required explicit modeling before estimating ST, LT and POP. A first example is repeated blood draws during the protocol, leading to what is called a “secular” trend, i.e. a progressive increase or decrease of protein abundance over time spent in protocol (see Supplementary Table 2). These proteins, including formimidoyltransferase cyclodeaminase (FTCD), and anthrax toxin receptor 2 (ANTXR2) are likely responsive to the experimental protocol itself (Extended Data Fig. 2a,b). As an example, ANTXR2 participates in extracellular matrix homeostasis and angiogenesis and is thus likely affected by repeated blood draws^79^. These effects were accounted for with a ‘cumulative hours’ fixed effect in the model, to ensure correct ST estimation (See Methods).

Another technical confounder identified in the data was hemolysis. In several samples, we observed abnormally high measured values for hemoglobin (HBB) over very short time intervals (Extended Data Fig. 2c). Although HBB can vary for biological reasons, including anemia, hemoglobinopathies, or severe dehydration, nearly 1,000-fold differences between samples collected two hours apart are more consistent with hemolysis during sample preprocessing. This interpretation was supported by review of the laboratory notes for the 17 most extreme plasma samples, among which 9 were annotated as hemolyzed (data not shown). Persistently high HBB levels could also reflect biological hemoglobin variation, including sickle-cell variants identified in the genetic analysis, and these two sources are difficult to distinguish in single-sample cohorts. We therefore examined whether other proteins showed the expected covariance with hemoglobin-related signals. As expected, Haptoglobin (HP) showed an inverse relationship with HBB, consistent with its role in binding free hemoglobin and promoting clearance of hemoglobin–haptoglobin complexes^80^ (Extended Data Fig. 2d). To account for both pre-analytical hemolysis and biological hemoglobin-related variation, we included hemoglobin level as a fixed effect in downstream models, thereby capturing additional proteins correlated with the hemolysis/hemoglobin axis that could have otherwise inflated ST estimates (Supplementary Table 2).

### The ST/LT/POP/GEN taxonomy characterizes plasma physiology

To assess biological significance of the metrics described above, we performed pathway enrichment analyses (restricted to proteins quantified in the SomaScan panel) to identify functional clusters of proteins associated with ST, LT, POP and GEN (Fig. 5a). All the identified pathways with respective adjusted P-values are available in Supplementary Table 4, and all the proteins displayed in the figure are listed in Supplementary Table 5. *Highly dynamic* (ST+/LT+) markers were associated with membrane trafficking pathways, receptor tyrosine kinase signaling and platelet activation and aggregation. These pathways capture processes responsive to both acute physiological fluctuations and slower systemic remodeling. As an example, the ‘Membrane Trafficking’ pathway is displayed at the individual protein level, including two proteins with multiple isoforms, Amyloid-beta precursor protein (APP) and Protein kinase B (PKB) (also called AKT, a serine/threonine kinase that serves as the central effector of the PI3K signaling pathway).

**Figure 5.**
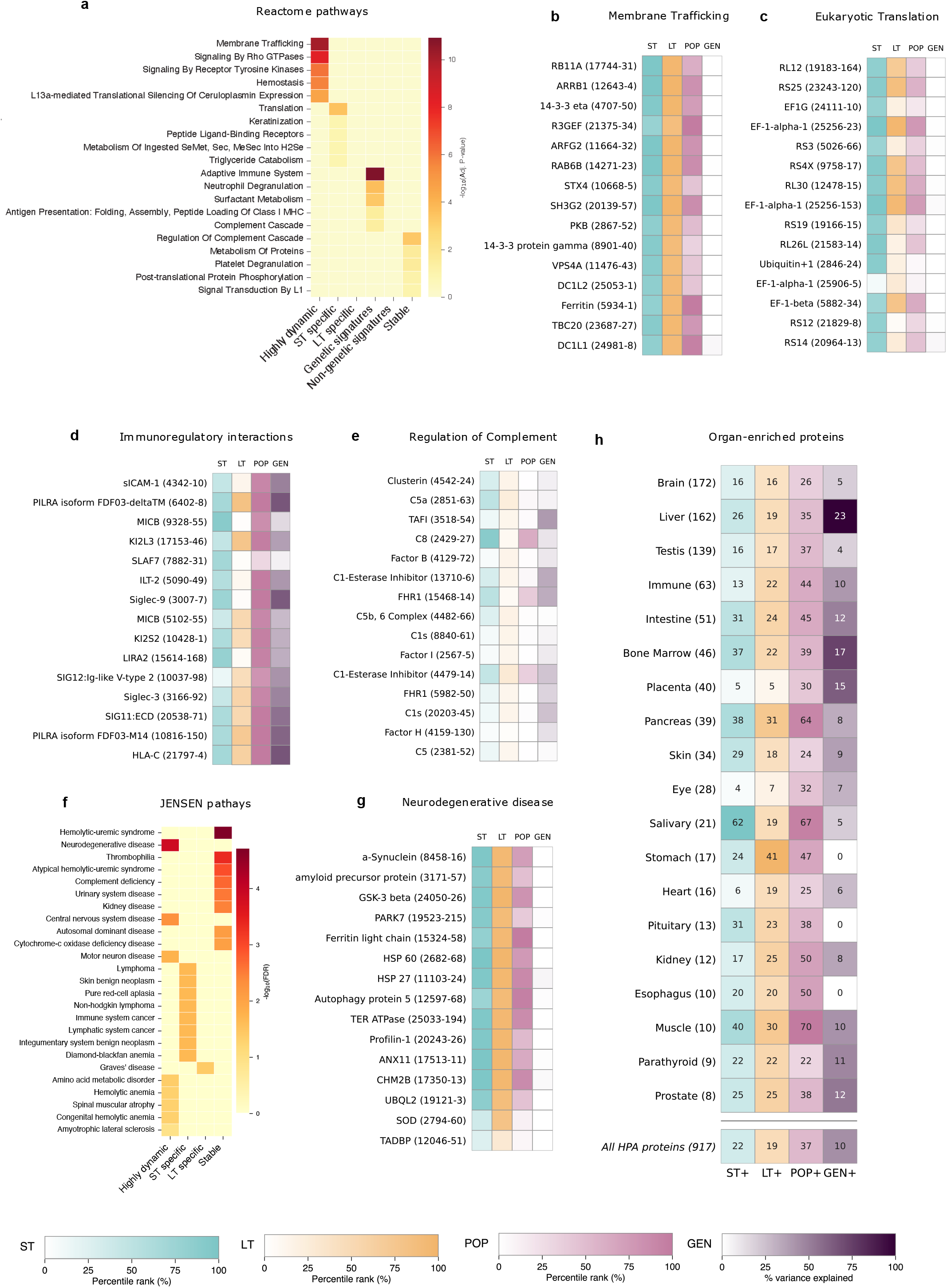
The variability-based taxonomy recapitulates plasma proteome physiology and disease. **a.** Hierarchical pathway analysis performed against the Reactome database. **b-e.** Heatmap of ST/LT/POP/GEN per aptamer, for the top pathways identified in a. **f.** Pathway analysis performed against JENSEN curated disease. **g.** Heatmap of ST/LT/POP/GEN per aptamer, for the top pathway identified in f. **h.** Heatmap of HPA organ-enriched proteins across variability categories. Values represent the percentage of proteins that are either ST+, LT+, POP+ or GEN+ in that specific organ, based on HPA definition. ‘Immune’ is comprised of lymphoid tissues. The last row indicates the percentage over all HPA-derived organ-enriched proteins. Category definitions: *Highly dynamic* = ST+/LT+; *ST+ specific* = ST+/¬LT+; *LT+ specific* = LT+/¬ST+; *Genetic signatures* = POP+/GEN+; *Non-genetic signatures* = POP+/¬GEN+; *Stable* = ST−/LT−/POP−. Categories are non-mutually exclusive (Extended Data Fig. 3a). (Definitions: ‘+’ = above upper quartile (Q75) ‘−’ = below lower quartile (Q25), ‘¬’ = exclusion; GEN+ = GEN > 20%). For pathway analysis, the custom background was set as the SomaScan 7K panel. Each aptamer is displayed with the naming convention ‘Protein Name (seq-ID)’. Row labelling indicates pathway name. Each pathway and associated proteins are available in Supplementary Tables 4 and 5. For heatmap figures, the legend indicates the rank of the protein in the ST/LT or POP category, for fair comparison independent of absolute value. *P*-values were adjusted by the Benjamini–Hochberg procedure. Terms were retained at adjusted P-value ≤ 0.15 with at least four overlapping proteins, then pruned by overlap coefficient ≥ 0.6. All the identified pathways with respective adjusted P-values are available in Supplementary Table 4. All the proteins belonging to pathways displayed in the figure are available in Supplementary Table 5. When the pathway identified contains more than 15 aptamers, a random selection of 15 aptamers was performed, except for neurodegenerative disease that is curated to also include ‘Motor neuron disease’.

Since *ST+* and *LT+* proteins largely overlap (Extended Data Fig. 3a) we used all 700 proteins that were ST+ but not LT+ to isolate short-term biological signatures (*ST-specific*, ST+/¬LT+). *ST+ specific* proteins were enriched in translation elongation and related ribosomal processes, in the cellular response to amino acid deficiency, and in selenoamino acid metabolism. Given their importance in xenobiotic processing, we also investigated proteins of the ADME class (Absorption, Distribution, Metabolism and Excretion) and found that these proteins showed high short-term variability (25/51 ST+; Extended Data Fig. 3b), consistent with short-timescale regulation of proteins involved in metabolism of drugs. As an example, the ‘Eukaryotic Translation Elongation’ pathway, comprising different elongation factors and ribosomal proteins, is displayed at the individual protein level (Fig. 5c). We also performed a mapping of proteins annotated as organ-specific in the HPA (Fig. 5h). Interestingly, pituitary was enriched in ST+ proteins, suggesting that diurnal hormonal secretion is captured in this category, along with salivary proteins that are more likely responsive to feeding schemes. In contrast, the same approach applied to the 700 LT+/¬ST+ proteins did not yield any significantly enriched pathway term, suggesting that *LT-specific* variation arises from more diverse mechanisms than the coherent short-term processes. Muscle-specific proteins, however, were found associated with LT+ variation, coherent with potential long-term physical activity variations.

As we expected, proteins with *genetic signature* (POP+/GEN+) were enriched for immune-related pathways, including ‘Immunoregulatory interactions between lymphoid and non-lymphoid cells’ (Fig. 5d) and components of the innate immune system. Additional enrichments included surfactant metabolism, and antigen presentation pathways. Furthermore, these proteins were annotated as enriched in immune-related organs such as the placenta and bone marrow (Fig. 5h). In contrast, proteins with *non-genetic signatures* (POP+/GEN-) did not yield any enriched pathway, suggesting a heterogeneous set of proteins for which variability is yet to be explained.

*Stable* proteins (ST–/LT–/POP–) yielded a coherent set of homeostatic pathways such as ‘Regulation of the complement cascade’ (Fig. 5e), ‘Platelet degranulation’, and ‘Insulin-like Growth Factor (IGF) transport and uptake by IGF binding proteins (IGFBPs)’. These *stable* pathways are coherent with our previous observation that highly abundant proteins in the plasma tend to have lower values of ST and LT. Nonetheless, complement component C8 (C8) still displayed ST+ variation (Fig. 5e). Similarly, several other components of the complement cascade were enriched in *genetic- signatures*, such as Complement factor H-related protein 2, 4 (CFHR2 and 4), Ficolin-2 (FCN2), mannose-binding protein C (MBL2) and vitronectin (VTN), all liver enriched (Fig. 5h). This suggests that selecting biomarkers for a given biological pathway must result from careful characterization of the variability profile of its members.

Next, we examined the enrichment for disease-associated gene sets (Curated from JENSEN^81,82^), and found that *highly dynamic* proteins were enriched for proteins involved in ‘Neurodegenerative disease’ (Fig. 5f), yielding a set of canonical and disease-linked neurodegeneration-associated proteins such as α-synuclein (SNCA), amyloid-beta precursor protein (APP), heat shock protein beta-1 (HSPB1), and annexin A11 (ANXA11) (Fig. 5g). This pattern is consistent with the known peripheral biology of several neurodegeneration-related proteins, including the marked abundance of SNCA in erythrocytes^83,84^ and the expression of multiple APP isoforms in circulating blood cells such as platelets and leukocytes^85^. Note that despite adjustment for hemolysis in the model, ST+ values persisted for SNCA, suggesting complex release. These findings suggest that plasma proteomics can capture blood-detectable neurodegeneration-associated signatures, while emphasizing the need to account for peripheral sources of variability when considering such proteins as biomarkers, for instance by using multiple assays or longitudinal measurements^86^.

We then asked whether proteins within the *genetic signature* category recapitulated known genotype–trait associations. Enrichment against curated Genome-wide Association Study (GWAS) Catalog^87^ terms showed that the *genetic signature* category captured proteins associated with blood cell traits, consistent with genetically determined variation in levels of circulating blood-related proteins such as the ABO groups (Extended Data Fig. 3c). This analysis also identified enrichment for the ‘Inflammatory bowel disease’ (IBD) GWAS term, indicating that some *genetic signature* proteins measurable in plasma are encoded by genes linked to IBD genetic risk. These proteins therefore connect disease-associated genetic architecture with genetically structured plasma abundance. If evaluated as biomarkers, their interpretation may benefit from genotype-aware modeling to distinguish inherited baseline differences from disease-associated or progression-related changes (Extended Data Fig. 3c,d).

## Discussion

Short-term variability is underappreciated because its study requires dense longitudinal sampling across multiple days and nights. By leveraging three longitudinal cohorts that encompass controlled day-to-day protocols and multi-year follow-up, this study jointly estimated ST, LT, POP, and GEN for 7,289 plasma protein targets. Mean ST was of the same order as LT (11.0% and 14.8% CV, respectively), while GEN explained an average of 4.0% of protein variance and was undetectable for 45% of probes. Short-term variation therefore affects at least as much of the plasma proteome as genetic variation: 1,822 probes were ST+ against 402 GEN+. This suggests that sparse longitudinal or cross-sectional designs may misattribute transient within-person variation to apparent LT or POP, and that ST should be considered alongside genetic background when designing and interpreting plasma proteomic studies. In our organ age-gap prediction case study, substantial fluctuations in predicted heart age coincided with high short-term variability of NT-proBNP, the model’s most heavily weighted predictor (ST = 26.2%; Fig. 1).

More generally, ST+ proteins pose a challenge for disease biomarker studies because their measured abundance can depend strongly on sampling context. When blood draw time, fasting status, sleep, posture or recent behavior differ between cases and controls, short-term physiological variation may either mimic disease-associated differences or obscure true disease effects. This issue is particularly relevant for diagnostic studies based on single blood samples, in which transient within-person variation cannot be distinguished from stable case-control differences. Standardized collection can reduce this confounding, whereas repeated sampling is required to estimate it directly^44^. This concern is illustrated by recently proposed candidates in studies where sampling time was not explicitly standardized. For instance, a recent study on tic disorders^88^ used a plasma ELISA assay to identify Chemokine (C-C motif) ligand 5 (CCL5) and Platelet-derived growth factors (PDGF-AB and PDGF-BB) as potential biomarkers. These proteins displayed substantial ST in our analysis, with values of 47.6%, 35.3% and 56.6%, respectively.

However, controlling for clock time cannot fully account for inter-individual differences in internal circadian phase, and circadian effects can only be distinguished from diurnal effects under controlled conditions. Nonetheless, it should still reduce variance introduced by diurnal behaviors, feeding, posture and sleep debt. In some studies, additional attention should be given to markers previously reported to have circadian effects. Four-jointed box protein 1 (FJX1), the most significant circadian protein in our previous study^50^, has been recently evaluated as a colorectal cancer marker, with moderate effect size and diagnostic performance^90^. Because cancer has been shown to disrupt an individual’s circadian rhythms^91,92^, controlling for clock time but not internal circadian phase might not be sufficient to account for circadian effects in such cases. Although we did not further partition ST into its drivers, future protocols could systematically manipulate these factors to isolate their effects.

Beyond ST confounding, we introduced a taxonomy derived from ST, LT, POP and GEN that complements recent studies of development, aging, tissue of origin and disease, which have characterized the expanding plasma proteome across both larger protein panels and broader cohort collections^5,33,94–96^. We show that our taxonomy links variability patterns to biological processes shaping protein trajectories. For instance, *highly dynamic* proteins were enriched in membrane trafficking and Rho GTPase signaling pathways, concordant with known diurnal variation in plasma extracellular vesicle size and secretion, and known diurnal liver proteome regulation^97,98^. *ADME* proteins showed higher short-term variability than the plasma proteome as a whole, coherent with chronopharmacology studies showing time-dependent (often circadian) regulation of xenobiotic metabolism^93,100,101^. As key determinants of drug metabolism, their marked ST may therefore be relevant, although plasma abundance reflects enzymatic activity only indirectly. Broadly used markers such as C-reactive protein (CRP) were also identified as ST+, suggesting that this marker is, as noted by others^99^, more dynamic than generally believed. Other *ST+* markers reflect rhythmic secretion rather than metabolism, such as TSH that is released in a diurnal and pulsatile manner^102^. On the other hand, *stable* proteins are enriched for complement, coagulation factors, and liver-associated proteins in line with the tight homeostatic regulation of abundant plasma proteins. Interestingly, individual proteins within the coagulation pathways displayed high variability, such as Plasminogen Activator Inhibitor-1 (PAI-1; ST+/LT+/POP+) which was previously linked to morning thrombosis peaks^104–107^. Finally, *genetic signature* proteins were enriched in immune pathways, recapitulating large pQTL studies showing strong genetic control of circulating immune proteins, a system characterized by inherent functional interindividual variation^5,95^.

Although the taxonomy showed coherent organization at the pathway level, individual proteins require assay- and proteoform-specific interpretation. For instance, we classified Fibronectin 1 (FN1) as ST+, whereas earlier plasma measurements reported relative stability over days, consistent with its long circulating half-life^111^. Given FN1’s extensive isoform diversity (plasma and cellular forms, with different fragments also quantified as ST+, see Supplementary Table 2), the SomaScan signal could in principle reflect a more variable fibronectin pool or probe-specific binding rather than total plasma fibronectin. However, the FN1 probe correlated well with Olink (Spearman ρ > 0.5)^53^, which argues against an assay-specific artifact and suggests its short-term variability is truly biological. More broadly, cross-platform concordance with Olink offers partial reassurance, but systematic comparisons show discrepancies can be substantial, probe-dependent, and biologically complex^5,112^. Interestingly, proteins with high correlation between the two platforms tended to be more frequently ST+, suggesting that these probes capture genuine short-term biological variability. On the other hand, poorly concordant probes may simply fail to track true signal and therefore appear artificially low in ST. Recent work further suggests that discordance between platforms partly arises from genetic effects influencing probe binding, potentially resulting in estimates of GEN that do not reflect genetic effects on biological protein abundance, but rather effects of genetic variants on assay efficacy^113^. As an example, we identified a protein-altering variant of alpha-2-HS-glycoprotein (AHSG) that is likely causing a disruption of aptamer binding (Supplementary Fig. 8).

Regarding modeling, our work proposes a framework that will help researchers prioritize protein candidates whose variability profiles are best aligned with the intended application. For instance, efforts to identify biomarkers of circadian phase or acute metabolic state should preferentially examine the ST+/LT−/POP− category, in which short-term responsiveness is coupled to limited long-term and interindividual variability; conversely, POP+/LT−/ST− proteins may provide stable individual baselines useful for risk stratification or diagnosis. The same framework should also inform the design of protein-specific studies: ST+ proteins may require larger sample sizes and/or repeated measures to detect disease-associated effects, consistent with variability-aware design principles^108,109^. LT+ proteins may be particularly suitable for monitoring disease progression, treatment response, or other longitudinal changes. GEN+ proteins may require genotype-aware interpretation to distinguish inherited baseline differences from disease-associated changes. Finally, these estimates could also be evaluated as priors in predictive models, for instance by testing whether down-weighting ST+ proteins improves transferability of single-sample discriminative models. The identification of highly abundant stable proteins in this dataset may also inform methodological choices for normalization techniques between batches, or controls for plasma assays, with multi-protein rather than single-protein normalization strategies^110^.

In conclusion, by jointly estimating ST, LT, POP, and GEN across 7,289 SomaScan protein targets, we define a structured taxonomy of plasma proteins and provide actionable priors for plasma characterization, biomarker selection, study design, and predictive models of health outcomes.

## Limitations of the study

Although our study reveals the importance of ST in study design and analysis, it does not necessarily reveal all possible ST effects and has limitations inherent to its design. For instance, secular proteins exhibiting protocol-related effects and proteins whose abundance can be altered by hemolysis (Supplementary Table 2) require explicit modeling, as was done here with cumulative hours spent in protocol and HBB levels. In addition, ST was estimated in a modest number of relatively young adult participants (n = 21) across two cohorts. Larger and more diverse cohorts will be required to determine how demographic and genetic factors influence ST, leading to more precise estimates. LT estimation was also limited to the ten-year follow-up available in MESA. Biological processes evolving over longer timescales or acquired during critical periods of development may therefore have been captured as POP at the expense of LT. Longer follow-up, with more diverse age ranges^29^ and precise mapping of exposure to disease will be needed to clearly distinguish persistent acquired set-points from reversible factors. The former may include developmental epigenetic marks, latent infections, or durable environmental exposures acquired at a specific time but stable thereafter. The latter may include modifiable states such as smoking, adiposity, disease activity, and medication exposure, which are more readily captured by our design. Regarding GEN, current estimates were derived from pQTLs discovered in MESA (n = 5,149) which includes diverse ancestries. These GEN estimates were highly correlated with those from a larger Icelandic SomaScan v4 cohort (deCODE, n = 35,559)^5^, which supports the robustness of protein-specific GEN estimates. However, some protein-specific genetic effects may have been underestimated at the current sample size, particularly for rare, ancestry-specific or modest-effect variants. Additionally, future work incorporating protein measures at multiple visits and times of the day will be needed to fully characterize how genetic variants impact measurement stability over time.

Finally, regarding platform specificity, we stress that variability estimates may differ across platforms because aptamer- and antibody-based assays capture distinct proteoforms, post-translational modifications, or protein complexes, thereby affecting binding and quantification. This highlights the need for protein-specific interpretation and complementary use of higher-resolution MS technology whenever feasible (although each platform holds its own set of limitations)^113–115^. Therefore, further work should test the transferability of our proposed variability taxonomy to different types of proteomics assays.

## STAR Methods

### Aims and overall design of the study

Our aim was to characterize short-term (ST), long-term (LT), population (POP) and genetic (GEN) variability of plasma proteins within a unified analytic framework. To determine the extent of short-term protein variation, we used two circadian-study related cohorts (S1, S2), and for long-term, population and genetics, a large-scale population cohort (MESA). SomaLogic was used for protein assessments. Plasma protein concentrations fluctuate due to a combination of technical noise and biological processes acting across multiple timescales. For each sample, the SomaScan v4 and v4.1 assays produced fluorescence-based abundance estimates for 4,985 and 7,289 proteins, respectively. To quantify these components and derive physiologically meaningful stability metrics, we modeled repeated SomaScan measurements using a hierarchical linear mixed-effects framework. SomaLogic provided intra-plate technical variances, and these were incorporated to separate measurement error from biological variation.

### Cohort descriptions

S1 and S2: Participants underwent continuous hourly blood sampling with every 2-hour plasma proteome assay in controlled laboratory conditions over 2-4 days to quantify short-term variability. For cohort S1, sampling began at waketime after a baseline 9-h in-laboratory sleep opportunity scheduled at each participant’s habitual times. Samples were collected across the 15-h wake episode and the following 9-h sleep opportunity. Upon awakening the next morning, participants began a 39-h Constant Routine, a protocol used to reveal circadian rhythms by controlling sleep-wake state, posture, fasting-feeding, lighting, and activity level. In this protocol, participants are restricted to wakeful bedrest in a semi-recumbent posture, continuous dim lighting, with hourly snacks equivalent to 1/24 of daily caloric and fluid needs. This study is followed by a recovery day and night in normal conditions. Cohort S2 began a 36-h Constant Routine upon awakening after a baseline 8-h night in the laboratory. Protocols for the SomaScan assay have been described previously^50^.

MESA exam 1, 4 and 5: The Multi-Ethnic Study of Atherosclerosis (MESA) is a study of the characteristics of subclinical cardiovascular disease (CVD) and the risk factors that predict progression to clinically overt CVD or progression of the subclinical disease^49^. MESA consists of a diverse, community-based sample of an initial 6,814 men and women aged 45-84 years without known CVD at baseline. Thirty-eight percent of the recruited participants were White, 28 percent African American, 22 percent Hispanic, and 12 percent of Chinese descent. Participants were recruited from six field centers across the United States: Baltimore City and Baltimore County, Maryland; Chicago, Illinois; Forsyth County, North Carolina; Los Angeles County, California; New York, New York; and St. Paul, Minnesota. The first examination took place over two years, from July 2000 to July 2002, and has been followed by additional examinations.

Participants are being followed for identification and characterization of CVD events, including acute myocardial infarction and other forms of coronary heart disease, stroke, and heart failure; for CVD interventions; and for mortality. Follow-up telephone interviews with participants or their proxies are attempted at least annually to identify new hospitalizations and diagnoses. MESA staff request information on hospital admissions for any reason, outpatient CVD diagnoses, and death. Death certificate records are obtained from state vital statistics or the National Death Index. MESA staff record International Classification of Disease (ICD) diagnosis codes for all hospitalizations and request medical records for those with ICD codes related to CVD. MESA staff also request outpatient records for potential CVD events. However, not all hospitalizations and/or CVD event records were fully documented. Protocols for blood sample collection and plasma extraction have been described at length elsewhere (https://internal.mesa-nhlbi.org/about/timelines-and-procedures).

### Ethics approval and consent to participate

The studies S1 and S2 were reviewed and approved by the Human Research Committee at Mass General Brigham (formerly Partners Healthcare) and were conducted in accordance with the principles outlined in the Declaration of Helsinki. Each participant provided written informed consent prior to taking part in the study. The use of the MESA (Multi-Ethnic Study of Atherosclerosis) cohort data has been approved by the MESA Coordinating Center to allow for accessing de-identified datasets through official NIH repositories. In addition, informed consent was obtained for extensive data sharing (dbGaP) and genetic/omic studies, including candidate genes (NHLBI CARe), genome-wide scans (NHLBI SHARe), exome sequencing (NHLBI ESP) and, most recently, the NHLBI TOPMed program. The entire study has also been reviewed and approved by Stanford University Institutional Review Boards.

### Organ-specific biological age

Organ age gaps were estimated with the published plasma-proteomic aging models of Oh et al.^56^ using their organage package, without re-training. Complete-protein matrices were normalized to the assay- version-matched reference (SomaScan v4.0 and v4.1 processed separately) to predict an age for each organ model, including Heart and Kidney. The organ age gap is the difference between the chronological age and the age predicted by the organ-specific age prediction model. Organ–protein assignments are taken directly from Oh et al^56^.

### Modeling of short term, long-term and population variability

Let *y_i_*_j*k*_ denote the log₁₀-transformed concentration for protein p, measured from subject i at long-term exam j and short-term sample k, collected at elapsed time *Δt_i_*_j*k*_ from the beginning of the continuous blood draw. To decompose variation across subjects, visits, and timepoints, we fitted the following mixed-effects model independently for each protein:

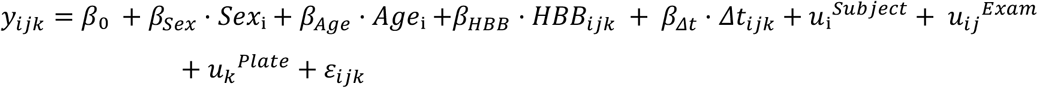

The model includes both fixed and random components to capture systematic and hierarchical sources of variation. Fixed effects consist of an intercept, sex, age, hemoglobin level (from SomaScan probe) and the elapsed sampling time Δt. A fixed-effect sex accounts for known baseline differences between male and female participants, while a fixed-effect slope coefficient associated with Δt models gradual physiological drift associated with prolonged continuous blood draw during the intensive sampling protocol (called “secular trend” discussed above).

Biological variation was modeled through subject- and exam-specific intercept random effects: a subject term *u_i_^Subject^* representing stable inter-individual differences in baseline protein abundance, and an exam term *u_ij_^Exam^* quantifying long-term biological drift across clinical visits separated by months to years. Technical batch effects were modeled via a plate-level intercept random effect *u_k_^Plate^* capturing differences arising from assay processing across plates and runs. The residual error term 3*_i_*_j*k*_encompassed short-term biological fluctuations occurring across hours to days, along with intra-plate technical variation intrinsic to each SOMAmer reagent. SomaLogic-provided assay-specific technical variances were incorporated into a version-dependent heteroscedastic residual structure, ensuring that short-term and technical noise were appropriately partitioned within the model. Variance components estimated on the log₁₀ scale were converted to percent coefficient of variation (%CV) using the standard transformation: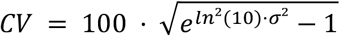

*Short-term biological variability* is extracted from the residual variance by subtracting the known SomaLogic intra-plate technical noise:

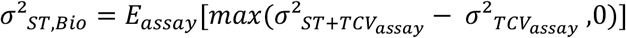

Assay-specific intra-plate technical variance for each protein and assay version was obtained directly from SomaLogic’s quality-control pipeline and treated as the minimal noise floor of the assay.

*Long-term biological variability* represents physiological change across visits, including the age effect. Because the exam-level variance component is already estimated net of the residual, no further subtraction of short-term or technical variance is applied:

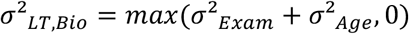

*Population-level variability* combines the subject-level random variance with the sex-related between-individual variance:

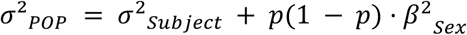

where p = 52% is the proportion of female participants. POP quantifies stable inter-individual differences that persist across short- and long-term timescales. Because the variance-to-CV transformation is exponential, %CV becomes very large for probes with extreme between-subject variance: 49 of 7,289 probes exceed POP = 100%, reflecting bimodal, genotype-driven signals rather than continuously distributed analytes.

Parameters were estimated using restricted maximum likelihood (REML) as implemented in the sommer package^116^. Uncertainty for fixed effects was quantified using Wald tests with two-sided p-values, and multiple testing across proteins was controlled using the Benjamini–Hochberg procedure. Variance components were estimated on the log-variance scale, and confidence intervals first computed as Wald intervals on this scale. The upper and lower bounds were then transformed to the percent coefficient-of-variation (CV) scale using the nonlinear variance-to-CV mapping, yielding confidence intervals by transformation of the Wald limits.

### pQTL mapping in MESA

Plasma samples from the MESA exam 1 were used for SomaScan 7K proteomics measurements. Whole genome sequence analysis was performed through TOPMed, using methods described previously^117^. To model protein measure variability expected due to genetic variation (GEN), we mapped protein quantitative trait loci (pQTL) in 5,149 contributing MESA participants with whole genome sequencing available through TOPMed and SomaScan 7K plasma protein measurements available at exam 1. Direct relatives, as determined by pedigree in MESA, and participants with flagged samples from SomaScan were excluded from analyses. When duplicate samples were present for a given individual, we kept only the TOPMed phase 2 sample.

Here, we used SomaScan protein measures with company-performed adaptive normalization by maximum likelihood (ANML) applied, according to SomaLogic recommendations. Protein measures were standardized and inverse normal transformed, then subsequently residualized with adjustment for covariates of age, biological sex (determined from WGS data), site, TOPMed ancestry principal components (PCs) 1-10, and protein measure PCs 1-10. The resulting residuals were inverse normal transformed and used as the outcome in pQTL mapping.

pQTL mapping was performed using the tensorQTL (version 1.0.10) *trans-* function^118^, with the adjustment for the same covariates protein measures were residualized upon, including variants with effect allele frequency (EAF) > 0.01 among the MESA participants. Loci to fine-map were selected using the EasyStrata INDEP function^119^, filtering for lead variants significant at P < 5x10^-8^ and at least 1 megabase (MB) apart. As long-range linkage disequilibrium (LD) can still exist at ranges greater than 1 Mb in some cases, we further calculated LD between lead variants using PLINK 1.90b3^120^ and filtered down to lead variants not in LD with one another to select loci to fine map (MESA r^2^ < 0.1).

To define genetic signals, we performed fine-mapping with SuSiE R (version 0.12.35)^121,122^ in 2Mb windows surrounding the lead variants identified previously, allowing for the number of credible sets per locus (i.e. SuSiE L parameter) to range from 1-10. For probes with at least 7 significant credible sets at a given locus in this initial run, we re-ran SuSiE at that locus, allowing the L parameter to range up to 20, as done in prior studies^123^. *cis*-pQTL were defined as credible sets within 1 Mb from the protein-encoding gene transcription start site (as defined primarily by GENCODE version 45^124^, incorporating RefSeq gene coordinates downloaded from the UCSC genome browser^125,126^ when no coordinates were available in GENCODE) containing at least one variant significant at p<5x10^-8^. *trans-* pQTL were defined as credible sets greater than 1 Mb from the protein-encoding gene transcription start site with at least one variant significant at p<6.8x10^-15^ (i.e. genome-wide significance corrected for the 7,275 proteins tested). We note that 14 proteins did not have gene coordinates available for the protein-encoding gene in the reference databases searched and were therefore excluded from pQTL results.

Furthermore, for a subset of probes, SuSiE generated credible sets with variants in linkage disequilibrium (LD). As these credible sets may not capture independent genetic signals, we calculated LD between variants in each *cis*-pQTL credible set for each protein or each *trans*-pQTL credible set. When at least 2 *cis-*pQTL or *trans-*pQTL credible sets contained variants in LD (defined here as r^2^>0.1), we filtered to keep only one of the credible sets, to ensure each credible set tags a distinct genetic signal, as performed previously^113^.

### Calculating variance explained by pQTL in MESA

For each protein with a significant pQTL credible set, we generated a polygenic score (PGS) in Plink version 1.90b3 ^120^ for the protein using the pQTL effect size (beta) for sentinel variants from each significant credible set as the weight. Notably, we excluded pQTL corresponding to SomaScan probe seq.19297.4, measuring glucose-6 phosphate-1-dehydrogenase (G6PD), due to evidence of substantial genomic inflation (λ=1.2) even after adjusting for 10 ancestry PCs in pQTL mapping. PGS were computed in the same set of MESA participants contributing to pQTL mapping. Variance explained by pQTL for each protein measure was calculated as the R^2^ value obtained when regressing the scaled PGS from inverse normal transformed protein measures in MESA. For the primary score, all significant *cis*- and/or *trans*-pQTL sentinel variants and corresponding variants were used. We note, for a small set of proteins, the *cis*-pQTL only score explained more protein measure variance than the combined *cis*- and *trans*-pQTL score, likely due to very long-range LD or ancestry differentiated variation resulting in non-random inheritance between some of the *cis* and *trans*-pQTL used in the score.

### Pathway and disease enrichment

Proteins were grouped into six non-mutually exclusive variability categories defined by panel-wide quartiles ST, LT, POP coefficients of variation, together with the estimated GEN (definitions in Fig. 5 legend). Over-representation analysis was performed for each category using gseapy.enrichr, a Python implementation/wrapper for Enrichr, with all measured plasma proteins used as background^127–129^ . Enrichments were queried against ‘Reactome_2022’, ‘Jensen DISEASES 2025’, and ‘GWAS Catalog 2025’^81,87,130^. Reactome 2022 is an expert-curated pathway knowledgebase, DISEASES integrates disease–gene associations from text mining and curated resources, and the NHGRI-EBI GWAS Catalog provides curated genotype–trait associations from published GWAS studies. *P*-values were adjusted by the Benjamini–Hochberg procedure. Terms were retained at adjusted *P* ≤ 0.15 with at least four overlapping proteins (min_genes = 4), then pruned by Szymkiewicz–Simpson overlap coefficient ≥ 0.6 (retaining the more significant pathway of each redundant pair); pure database identifiers were excluded. Cross-category heatmaps aggregate the top five redundancy-reduced terms per category.

### Per-aptamer visualisation

For each variability category, the most significant Reactome pathway retained after redundancy reduction was selected, and its constituent proteins were displayed as a square-cell heatmap with one row per protein and four columns corresponding to ST, LT, POP and GEN. Cell colour for ST, LT and POP was determined by the global percentile rank of the corresponding CV value, using method=’min’ so that proteins with CV = 0 collapsed at the white end of the scale. GEN was displayed on its native scale, as percentage variance explained, because of its zero-inflated distribution. The same heatmap format was applied to a curated neurodegenerative-disease panel and to the GWAS-derived IBD panel.

### Drug-target distribution

Aptamers mapping to the Reactome ‘DRUG_ADME’ gene set were intersected with the measured plasma-protein panel. Per-gene variability was then displayed using the same four-column heatmap, allowing the distribution of ADME-related drug targets across the ST, LT, POP and GEN axes to be read directly.

### Tissue-specificity enrichment

Tissue assignments were obtained from the HPA and intersected with each variability category. HPA tissue annotations are based on integrated transcriptomic and antibody-based proteomic profiling across human tissues and organs^57^. For each organ × variability pair, the percentage of proteins in the upper quartile (or GEN > 20%) in that organ was reported compared to the total number of proteins in that organ.

## Resource Availability

### Data availability

Constant Routine material (samples) used in the study is limited in quantity, but requests will be considered on a case-by-case basis. Individual proteomic and genetic data are considered identifiable, but all summary statistics are available in supplementary material. Access for data use from MESA needs pre-approval for a specific use through the MESA Coordinating Center. A full list of participating MESA investigators and institutions can be found at http://www.mesa-nhlbi.org

### Code availability

All analyses were performed using our repository stable-proteins available online at time of publication.

## Acknowledgments

Proteomic measurements and analysis of the data have been funded by gifts to E. Mignot, a contract from Takeda Pharmaceuticals, and by NIH US National Institutes of Health grant R01HL148704. Collection of the samples from study S1 was supported by a grant from the US National Institutes of Health [R01 HL148704]. Collection of the samples from study S2 was supported by a grant from the Office of Naval Research [N00014-15-1-2408]. Studies S1 and S2 were carried out at the Brigham and Women’s Hospital Center for Clinical Investigation, with support from Harvard Catalyst, The Harvard Clinical and Translational Science Center [National Center for Advancing Translational Sciences, NIH Award UL1 TR002541] and financial contributions from Brigham and Women’s Hospital, Harvard University, and its affiliated academic healthcare centers. Whole genome sequencing (WGS) for the Trans-Omics in Precision Medicine (TOPMed) program was supported by the National Heart, Lung and Blood Institute (NHLBI). WGS for “NHLBI TOPMed: Multi-Ethnic Study of Atherosclerosis (MESA)” (phs001416.v1.p1) was performed at the Broad Institute of MIT and Harvard (3U54HG003067-13S1). Centralized read mapping and genotype calling, along with variant quality metrics and filtering were provided by the TOPMed Informatics Research Center (3R01HL-117626- 02S1). Phenotype harmonization, data management, sample-identity QC, and general study coordination, were provided by the TOPMed Data Coordinating Center (3R01HL-120393-02S1). MESA and the MESA SHARe project are conducted and supported by the National Heart, Lung, and Blood Institute (NHLBI) in collaboration with MESA investigators. Support for MESA is provided by contracts 75N92025D00022, 75N92020D00001, HHSN268201500003I, N01-HC-95159, 75N92025D00026, 75N92020D00005, N01-HC-95160, 75N92020D00002, N01-HC-95161, 75N92025D00024, 75N92020D00003, N01-HC-95162, 75N92025D00027, 75N92020D00006, N01- HC-95163, 75N92025D00025, 75N92020D00004, N01-HC-95164, 75N92025D00028, 75N92020D00007, N01-HC-95165, N01-HC-95166, N01-HC-95167, N01-HC-95168, N01-HC- 95169, UL1-TR-000040, UL1-TR-001079, UL1-TR-001420, UL1TR001881, and R01HL105756.

We would like to thank the technical staff members of the BWH Division of Sleep and Circadian Medicine who assisted with the participant recruitment, screening, and study execution for cohorts S1 and S2; Enmanuel Pardilla-Delgado, Arturo Arrona-Palacios, and Noelia Ruiz-Herrera for overseeing the data collection segments of studies S1 and S2; Ms. Audra S. Murphy and Dr. Jing Zhang for coordinating the sample inventory, sorting, and shipment; and the technical, nursing, and dietary staff of the BWH Center for Clinical Investigation for assisting with the sample collection and processing. The authors also would like to thank the MESA participants and the MESA investigators and staff for their valuable contributions. Finally, we also wish to thank Matthew Weaver, Eric Yu, Camille Chataing, and Aaron Robinson for advice and suggestions on the manuscript.

## Author contributions

D.B., A.S.: Conceptualization, Statistical Analysis, Writing, Review & Editing; J.C.N., M.G.: Genetics, Statistical Analysis, Writing - Review & Editing; R.G.: Figures, Writing, Review & Editing; R.Du., P.G., J.I.R., K.D.T., S.S.R., P.Y.L., A.C.W., M.Y.M., R.De., K.-M.Z.: Data acquisition, Review & Editing; L.M.R.: Supervision, Writing - Review & Editing; C.A.C., J.F.D.: Supervision, Funding Acquisition, Data acquisition, Review & Editing; E.M.: Project Administration, Funding Acquisition, Conceptualization, Supervision, Writing, Review & Editing.

All authors have reviewed and approved the final version of the manuscript.

---

**Extended Data Figure 1.**
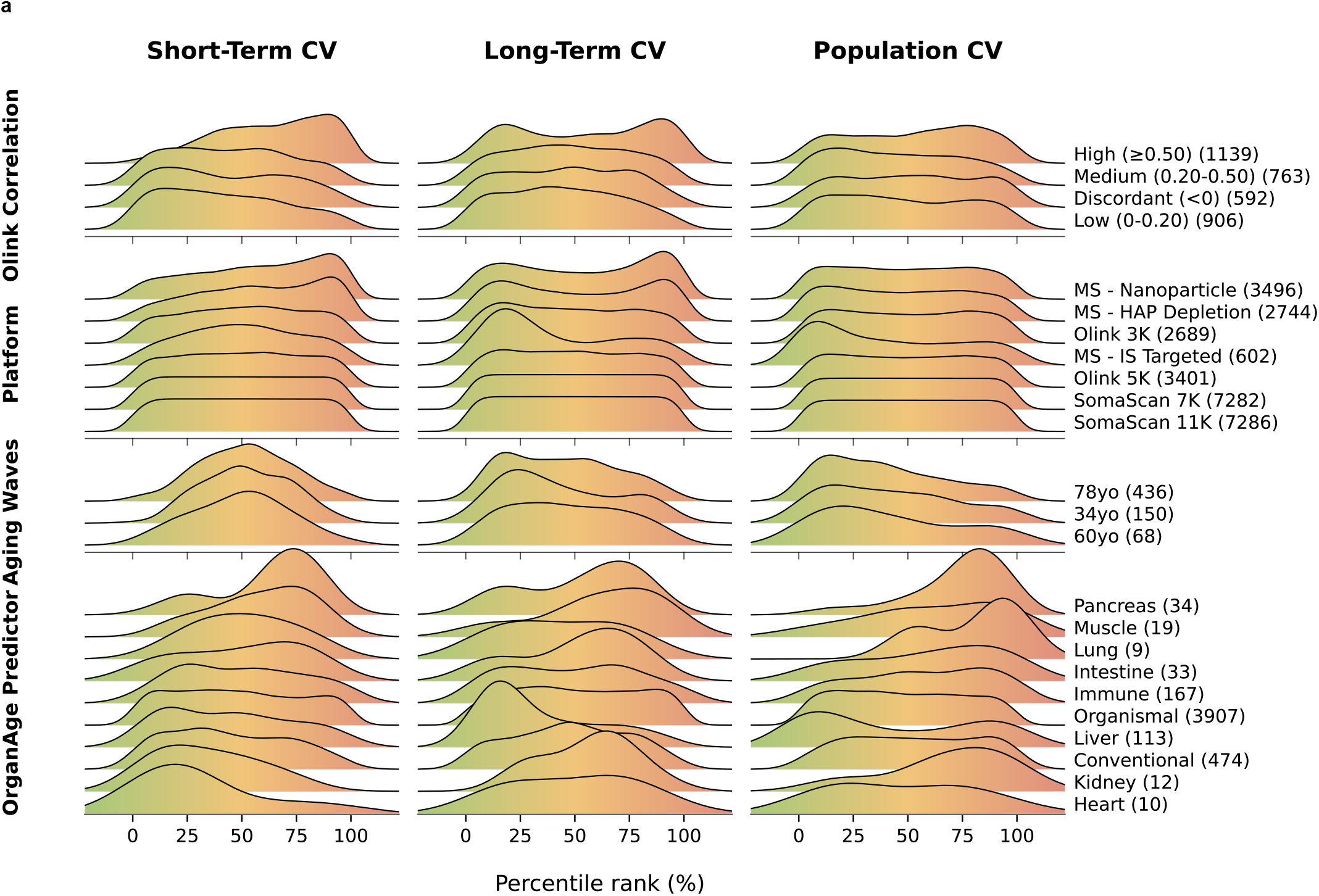
Supplementary characterization of ST, LT, POP and GEN. **a.** Ridgeline plots show the percentile-ranked variability of 7,289 SomaScan protein targets for short-term variability (ST, first column), long-term variability (LT, middle column), and inter-individual variability (POP, last column). Rows correspond to four annotation groups: agreement with Olink measurements (discordant, low, medium, high correlation), analytical platform (mass spectrometry (MS), MS with depletion, nanoparticle MS, Olink 3K, Olink 5K, SomaScan 7K, SomaScan 11K), aging-wave responsiveness^31^ (34-year, 60-year, 78-year groups), and predictors of organ age^56^ (heart, kidney, lung, liver, conventional plasma, organismal, immune, intestine, muscle, pancreas). Background color bands provide interpretive ranges for each metric (green: low variability, yellow: moderate, red: high).

**Extended Data Figure 2.**
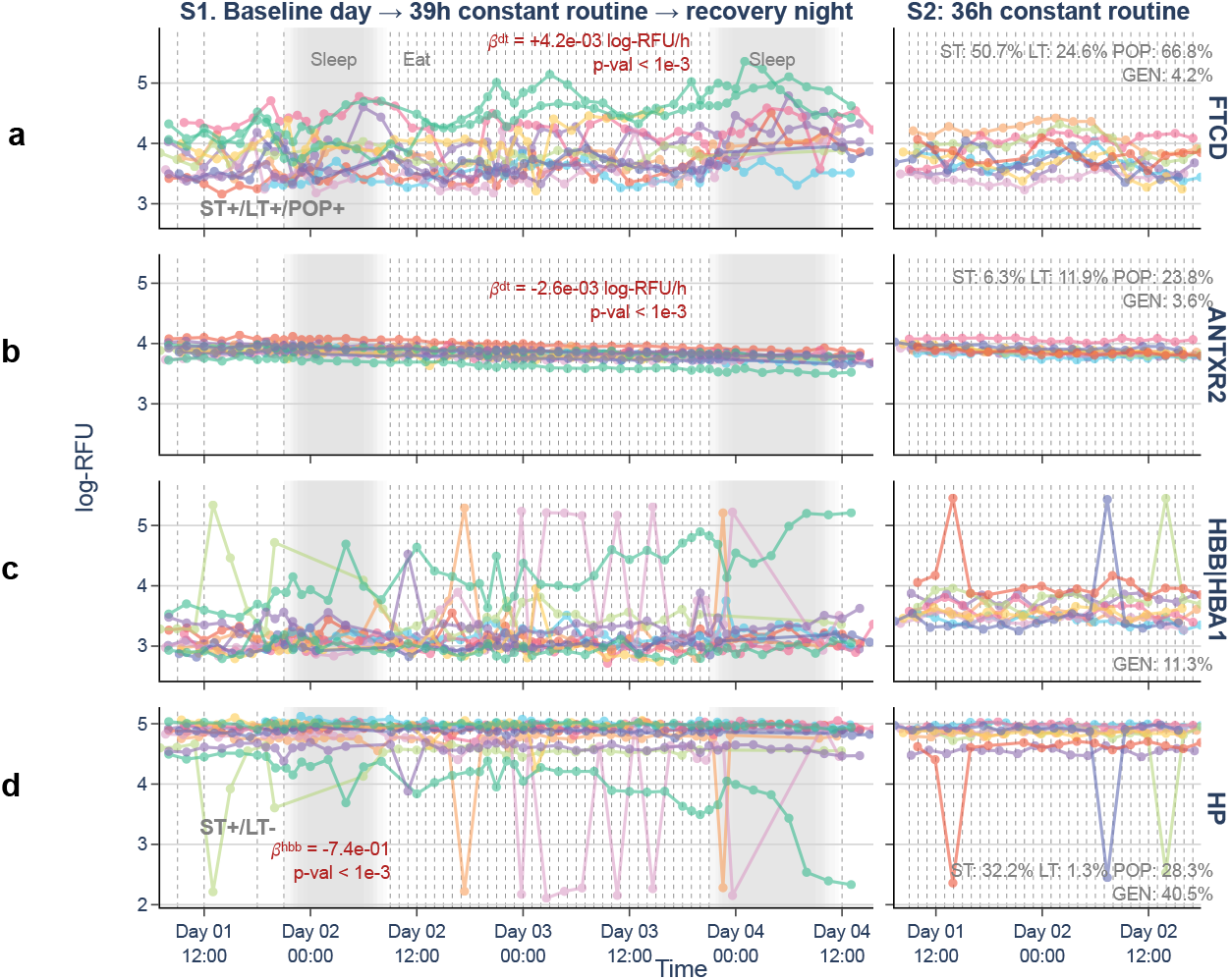
Representative protein trajectories subject to protocol-related secular drift and hemolysis. Representative protein trajectories exhibiting secular drift and hemolysis during Constant Routine protocols. Several proteins were identified as drifting during the longitudinal protocols. **a.** Representative trajectory of FTCD, with increasing secular drift imposed on ST+ variation. **b.** Representative trajectory of ANTXR2, with decreasing secular drift on an otherwise stable protein. Note that in the case of ANTXR2 there is no superimposed ST variation. In log-scale the drift is moderate but consistent across participants. **c.** Representative trajectory of HBB, displaying either longitudinally increasing hemolysis (turquoise participant) or hemolyzed samples (spikes). **d.** Representative trajectory of HP that antagonizes free HBB, thus inversely varying with HBB. β log-RFU/h is indicated in red along with P value. For each protein, longitudinal plasma abundances (log-RFU) are displayed for S1 and S2 in the first two columns, with participants indicated by color. For both cohorts, the x-axis indicates the day and hour of the sampling protocol; shaded regions denote scheduled sleep opportunities, and vertical dashed lines indicate meals during baseline and recovery days and snacks during the Constant Routine protocol. S1 (n = 12; 518 samples) was collected across a baseline day, a 39-h Constant Routine protocol, and a recovery night, whereas S2 (n = 9; 152 samples) was collected during a 36-h Constant Routine protocol.

**Extended Data Figure 3.**
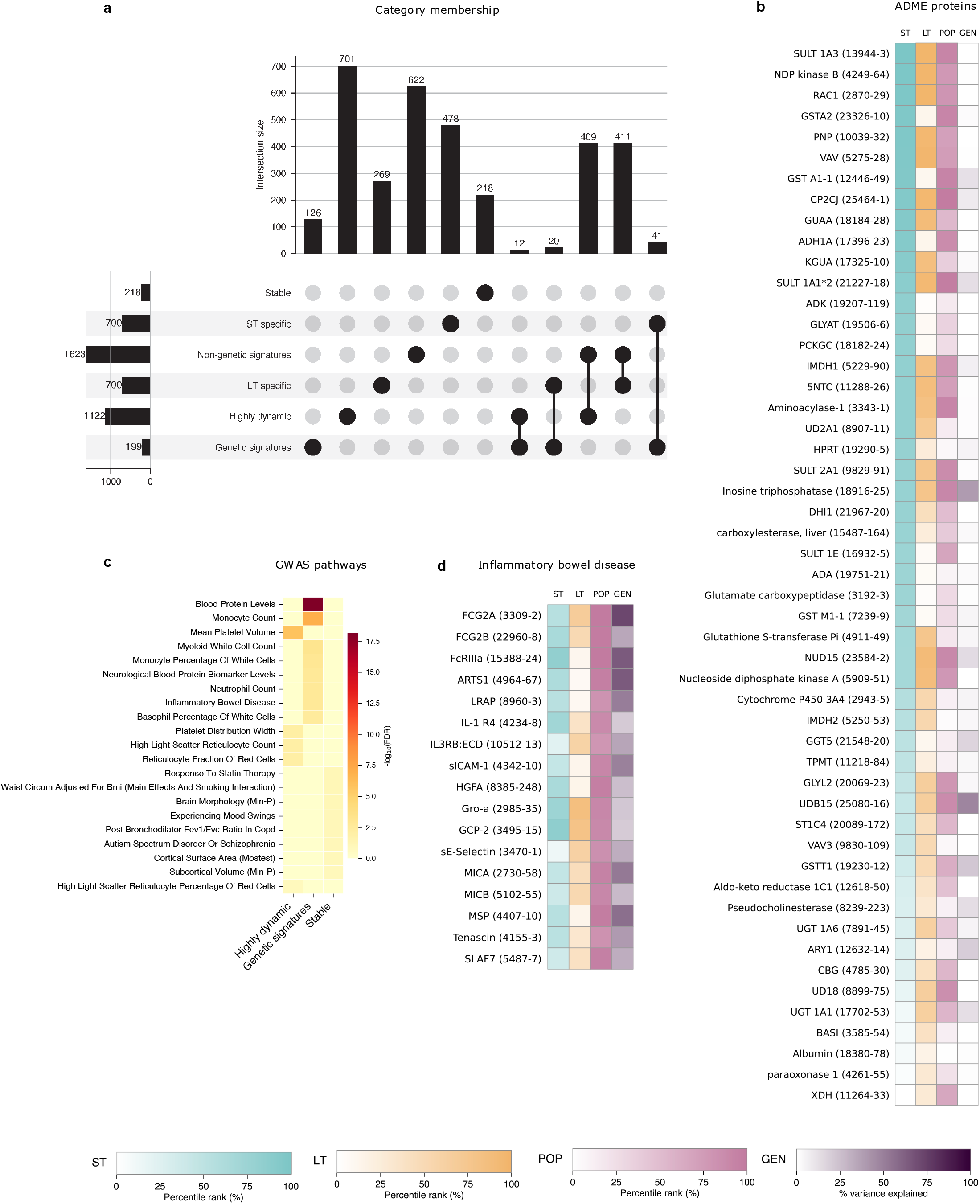
The variability-based taxonomy recapitulates plasma proteome physiology and disease. **a.** Upset plot depicting the numbers of proteins belonging to each of the protein categories analyzed by pathway enrichment. Total size of the category is indicated on the left horizontal bar plots (from 199 to 1623). Memberships and overlap are indicated by the dots and corresponding vertical bar plots on the top. As an example, *Genetic signatures* contain 199 aptamers in total, with 126 only in that category, 12 overlapping with *Highly dynamic*, 41 with *ST+ specific*, and 20 with *LT+ specific*. **b.** Heatmap of ST/LT/POP/GEN for the 51 absorption, distribution, metabolism, and excretion (ADME) proteins present in the SomaScan 7K panel. **c.** GWAS-enriched hits across defined categories. GWAS associated proteins are extracted from the NHGRI-EBI GWAS Catalog^87^. **d.** Heatmap of ST/LT/POP/GEN per aptamer, for ‘Inflammatory bowel disease’, identified in c. Categories: *Highly dynamic* = ST+/LT+; *ST+ specific* = ST+/¬LT+; *LT+ specific* = LT+/¬ST+; *Genetic signatures* = POP+/GEN+; *Non-genetic signatures* = POP+/¬GEN+; *Stable* = ST−/LT−/POP−. Categories are non-mutually exclusive (Extended Data Fig. 3a). (Definitions: ‘+’ = above upper quartile (Q75) ‘−’ = below lower quartile (Q25), ‘¬’ = exclusion; GEN+ = GEN > 20%). For pathway analysis, the custom background was set as the SomaScan 7K panel. Each aptamer is displayed with the naming convention ‘Protein Name (seq-ID)’. Row labelling indicates pathway name. Each pathway and associated proteins are available in Supplementary Tables 4 and 5. For heatmap figures, the legend indicates the rank of the protein in the ST/LT or POP category, for fair comparison independent of absolute value. *P*-values were adjusted by the Benjamini–Hochberg procedure. Terms were retained at adjusted *P* ≤ 0.15 with at least four overlapping proteins, then pruned by overlap coefficient ≥ 0.6.

## References

1. Palstrøm, N. B., Matthiesen, R., Rasmussen, L. M. & Beck, H. C. Recent Developments in Clinical Plasma Proteomics—Applied to Cardiovascular Research. Biomedicines 10, 162 (2022).

2. Mardinoglu, A. et al. Longitudinal big biological data in the AI era. Mol. Syst. Biol. 21, 1147–1165 (2025).

3. Ganz, P. et al. Development and Validation of a Protein-Based Risk Score for Cardiovascular Outcomes Among Patients With Stable Coronary Heart Disease. JAMA 315, 2532–2541 (2016).

4. Shi, L. et al. Discovery and validation of plasma proteomic biomarkers relating to brain amyloid burden by SOMAscan assay. Alzheimer’s Dement. 15, 1478–1488 (2019).

5. Eldjarn, G. H. et al. Large-scale plasma proteomics comparisons through genetics and disease associations. Nature 622, 348–358 (2023).

6. Kolasa, K. et al. Systematic reviews of machine learning in healthcare: a literature review. Expert Rev. Pharmacoeconomics Outcomes Res. 24, 63–115 (2024).

7. Al-Tashi, Q. et al. Machine Learning Models for the Identification of Prognostic and Predictive Cancer Biomarkers: A Systematic Review. Int. J. Mol. Sci. 24, 7781 (2023).

8. Winchester, L. et al. Identification of a possible proteomic biomarker in Parkinson’s disease: discovery and replication in blood, brain and cerebrospinal fluid. Brain Commun. 5, fcac343 (2022).

9. Prelaj, A. et al. Artificial intelligence for predictive biomarker discovery in immuno-oncology: a systematic review. Ann. Oncol. 35, 29–65 (2024).

10. Blanco, K. et al. Systematic review: fluid biomarkers and machine learning methods to improve the diagnosis from mild cognitive impairment to Alzheimer’s disease. Alzheimer’s Res. Ther. 15, 176 (2023).

11. Ng, S., Masarone, S., Watson, D. & Barnes, M. R. The benefits and pitfalls of machine learning for biomarker discovery. Cell Tissue Res. 394, 17–31 (2023).

12. Xing, X. et al. Proteomics-driven noninvasive screening of circulating serum protein panels for the early diagnosis of hepatocellular carcinoma. Nat. Commun. 14, 8392 (2023).

13. Dregoesc, M. I. et al. Relation Between Plasma Proteomics Analysis and Major Adverse Cardiovascular Events in Patients With Stable Coronary Artery Disease. Front. Cardiovasc. Med. 9, 731325 (2022).

14. Schuermans, A. et al. Integrative proteomic analyses across common cardiac diseases yield mechanistic insights and enhanced prediction. *Nat*. Cardiovasc. Res. 3, 1516–1530 (2024).

15. Dodig-Crnković, T. et al. Facets of individual-specific health signatures determined from longitudinal plasma proteome profiling. EBioMedicine 57, 102854 (2020).

16. Joshi, A. & Mayr, M. In Aptamers They Trust. Circulation 138, 2482–2485 (2018).

17. Candia, J., Daya, G. N., Tanaka, T., Ferrucci, L. & Walker, K. A. Assessment of variability in the plasma 7K SomaScan proteomics assay. Scientific Reports 12, 17147 (2022).

18. Haslam, D. E. et al. Stability and reproducibility of proteomic profiles in epidemiological studies: comparing the Olink and SOMAscan platforms. PROTEOMICS 22, e2100170 (2022).

19. Candia, J. et al. Variability of 7K and 11K SomaScan Plasma Proteomics Assays. J. Proteome Res. 23, 5531–5539 (2024).

20. Carlyle, B. C. et al. Technical Performance Evaluation of Olink Proximity Extension Assay for Blood-Based Biomarker Discovery in Longitudinal Studies of Alzheimer’s Disease. Front. Neurol. 13, 889647 (2022).

21. Kirsher, D. Y. et al. The Current Landscape of Plasma Proteomics: Technical Advances, Biological Insights, and Biomarker Discovery. bioRxiv 2025.02.14.638375 (2025) doi:10.1101/2025.02.14.638375.

22. Wang, B. et al. Comparative studies of 2168 plasma proteins measured by two affinity-based platforms in 4000 Chinese adults. Nat. Commun. 16, 1869 (2025).

23. Koprulu, M. et al. Sex differences in the genetic regulation of the human plasma proteome. Nat. Commun. 16, 4001 (2025).

24. Carrasco-Zanini, J. et al. Mapping biological influences on the human plasma proteome beyond the genome. Nat. Metab. 6, 2010–2023 (2024).

25. Kim, C. H. et al. Stability and reproducibility of proteomic profiles measured with an aptamer-based platform. Sci. Rep. 8, 8382 (2018).

26. Núñez, E. et al. Unbiased plasma proteomics discovery of biomarkers for improved detection of subclinical atherosclerosis. eBioMedicine 76, 103874 (2022).

27. Vasaikar, S. V. et al. A comprehensive platform for analyzing longitudinal multi-omics data. Nat. Commun. 14, 1684 (2023).

28. Lakshmikanth, T. et al. Human Immune System Variation during 1 Year. Cell Rep. 32, 107923 (2020).

29. Ingvarsdottir, H. K. et al. Long-term temporal stability of circulating proteins in older adults. Nat. Commun. (2026) doi:10.1038/s41467-026-72957-w.

30. Tanaka, T. et al. Plasma proteomic biomarker signature of age predicts health and life span. eLife 9, e61073 (2020).

31. Lehallier, B. et al. Undulating changes in human plasma proteome profiles across the lifespan. Nat. Med. 25, 1843–1850 (2019).

32. Rao, Z. et al. A Novel Longitudinal Proteomic Aging Index Predicts Mortality, Multimorbidity, and Frailty in Older Adults. Aging Cell 25, e70317 (2026).

33. Álvez, M. B. et al. A human pan-disease blood atlas of the circulating proteome. Science eadx2678 (2025) doi:10.1126/science.adx2678.

34. Bergström, S. et al. Longitudinal protein profiling of blood during childhood into early adulthood. Nat. Commun. 17, 3700 (2026).

35. Pietzner, M. et al. Systemic proteome adaptions to 7-day complete caloric restriction in humans. Nat. Metab. 6, 764–777 (2024).

36. Mi, M. Y. et al. Plasma Proteomic Kinetics in Response to Acute Exercise. Mol. Cell. Proteom. 22, 100601 (2023).

37. Brandão, L. E. M. et al. The overlooked trio: sleep duration, sampling time and physical exercise alter levels of olink-assessed blood biomarkers of cardiovascular risk. Biomark. Res. 13, 67 (2025).

38. Lee-Ødegård, S. et al. Serum proteomic profiling of physical activity reveals CD300LG as a novel exerkine with a potential causal link to glucose homeostasis. eLife 13, RP96535 (2024).

39. Robbins, J. M., et al. Plasma proteomic changes in response to exercise training are associated with cardiorespiratory fitness adaptations. JCI Insight 8, e165867 (2023).

40. Kreft, I. C. et al. Mass spectrometry-based analysis on the impact of whole blood donation on the global plasma proteome. Transfusion 63, 564–573 (2023).

41. Lotfi, R., Kroll, C., Plonné, D., Jahrsdörfer, B. & Schrezenmeier, H. Hepcidin/Ferritin Quotient Helps to Predict Spontaneous Recovery from Iron Loss following Blood Donation. Transfus. Med. Hemotherapy 42, 390–395 (2015).

42. Moazzen, S. et al. Ferritin Trajectories over Repeated Whole Blood Donations: Results from the FIND+ Study. J. Clin. Med. 11, 3581 (2022).

43. Deota, S. et al. The time is now: accounting for time-of-day effects to improve reproducibility and translation of metabolism research. Nat. Metab. 7, 454–468 (2025).

44. Spick, M. et al. Challenges and opportunities for statistical power and biomarker identification arising from rhythmic variation in proteomics. *npj Biol*. Timing Sleep 2, 3 (2025).

45. Wang, W. et al. Desynchronizing the sleep--–wake cycle from circadian timing to assess their separate contributions to physiology and behaviour and to estimate intrinsic circadian period. Nat. Protoc. 18, 579–603 (2023).

46. Duffy, J. F. & Dijk, D.-J. Getting Through to Circadian Oscillators: Why Use Constant Routines? J. Biol. Rhythm. 17, 4–13 (2002).

47. Dijk, D.-J. & Duffy, J. F. Novel Approaches for Assessing Circadian Rhythmicity in Humans: A Review. J. Biol. Rhythm. 35, 421–438 (2020).

48. Vaquer-Alicea, A., Yu, J., Liu, H. & Lucey, B. P. Plasma and cerebrospinal fluid proteomic signatures of acutely sleep-deprived humans: an exploratory study. Sleep Adv. 4, zpad047 (2023).

49. Bild, D. E. et al. Multi-Ethnic Study of Atherosclerosis: Objectives and Design. Am. J. Epidemiology 156, 871–881 (2002).

50. Specht, A. et al. Circadian protein expression patterns in healthy young adults. Sleep Health 10, S41–S51 (2024).

51. Parcha, V. et al. Chronobiology of Natriuretic Peptides and Blood Pressure in Lean and Obese Individuals. J. Am. Coll. Cardiol. 77, 2291–2303 (2021).

52. Young, M. E. et al. A brief history of circadian time in the heart. J. Mol. Cell. Cardiol. 207, 66–80 (2025).

53. Kirsher, D. Y. et al. Current landscape of plasma proteomics from technical innovations to biological insights and biomarker discovery. Commun. Chem. 8, 279 (2025).

54. Salomon, R. M. et al. Diurnal variation of cerebrospinal fluid hypocretin-1 (Orexin-A) levels in control and depressed subjects. Biol. Psychiatry 54, 96–104 (2003).

55. Mignot, E. et al. The Role of Cerebrospinal Fluid Hypocretin Measurement in the Diagnosis of Narcolepsy and Other Hypersomnias. Arch. Neurol. 59, 1553–1562 (2002).

56. Oh, H. S.-H. et al. Organ aging signatures in the plasma proteome track health and disease. Nature 624, 164–172 (2023).

57. Uhlén, M. et al. Tissue-based map of the human proteome. Science 347, 1260419 (2015).

58. Sun, B. B. et al. Genomic atlas of the human plasma proteome. Nature 558, 73–79 (2018).

59. Moore, C. et al. The INTERVAL trial to determine whether intervals between blood donations can be safely and acceptably decreased to optimise blood supply: study protocol for a randomised controlled trial. Trials 15, 363 (2014).

60. Wang, X. Chapter Three Pleiotrophin: Activity and mechanism. Adv. Clin. Chem. 98, 51–89 (2020).

61. Vandenberghe-Dürr, S., Gilliet, M. & Domizio, J. D. OLFM4 regulates the antimicrobial and DNA binding activity of neutrophil cationic proteins. Cell Rep. 43, 114863 (2024).

62. Clemmensen, S. N. et al. Plasma levels of OLFM4 in normals and patients with gastrointestinal cancer. J. Cell. Mol. Med. 19, 2865–2873 (2015).

63. Alder, M. N., Opoka, A. M., Lahni, P., Hildeman, D. A. & Wong, H. R. Olfactomedin-4 Is a Candidate Marker for a Pathogenic Neutrophil Subset in Septic Shock. Crit. Care Med. 45, e426– e432 (2017).

64. Liu, W. & Rodgers, G. P. Olfactomedin 4 Is a Biomarker for the Severity of Infectious Diseases. Open Forum Infect. Dis. 9, ofac061 (2022).

65. Lu, H. et al. The BRCA2-Interacting Protein BCCIP Functions in RAD51 and BRCA2 Focus Formation and Homologous Recombinational Repair. Mol. Cell. Biol. 25, 1949–1957 (2005).

66. Ye, C. et al. BCCIP is required for nucleolar recruitment of eIF6 and 12S pre-rRNA production during 60S ribosome biogenesis. Nucleic Acids Res. 48, gkaa1114- (2020).

67. Le, H. P., Heyer, W.-D. & Liu, J. Guardians of the Genome: BRCA2 and Its Partners. Genes 12, 1229 (2021).

68. Tang, V. T., et al. Functional overlap between the mammalian Sar1a and Sar1b paralogs in vivo. Proc. Natl. Acad. Sci. 121, e2322164121 (2024).

69. Petrosyan, A., Cheng, P.-W., Clemens, D. L. & Casey, C. A. Downregulation of the small GTPase SAR1A: a key event underlying alcohol-induced Golgi fragmentation in hepatocytes. Sci. Rep. 5, 17127 (2015).

70. Sempere, V. P. et al. HLA and KIR genetic association and NK cells in anti-NMDAR encephalitis. Front. Immunol. 15, 1423149 (2024).

71. Trowsdale, J. et al. The genomic context of natural killer receptor extended gene families. Immunol. Rev. 181, 20–38 (2001).

72. Traherne, J. A. et al. Mechanisms of copy number variation and hybrid gene formation in the KIR immune gene complex. Hum. Mol. Genet. 19, 737–751 (2010).

73. Agrawal, S. & Prakash, S. Significance of KIR like natural killer cell receptors in autoimmune disorders. Clin. Immunol. 216, 108449 (2020).

74. Amorim, L. M. et al. High-Resolution Characterization of KIR Genes in a Large North American Cohort Reveals Novel Details of Structural and Sequence Diversity. Front. Immunol. 12, 674778 (2021).

75. Oussalah, A., Levy, J., Filhine-Trésarrieu, P., Namour, F. & Guéant, J.-L. Association of TCN2 rs1801198 c.776G>C polymorphism with markers of one-carbon metabolism and related diseases: a systematic review and meta-analysis of genetic association studies. Am. J. Clin. Nutr. **106**, 1142–1156 (2017).

76. Farinas, A. et al. Disruption of the cerebrospinal fluid–plasma protein balance in cognitive impairment and aging. Nat. Med. 31, 2578–2589 (2025).

77. Lohr, N. J. et al. Human ITCH E3 Ubiquitin Ligase Deficiency Causes Syndromic Multisystem Autoimmune Disease. Am. J. Hum. Genet. 86, 447–453 (2010).

78. Moser, E. K. & Oliver, P. M. Regulation of autoimmune disease by the E3 ubiquitin ligase Itch. Cell. Immunol. 340, 103916 (2019).

79. Sergeeva, O. A. & Goot, F. G. van der. Converging physiological roles of the anthrax toxin receptors. F1000Research **8**, F1000 Faculty Rev-1415 (2019).

80. Schaer, D. J., Vinchi, F., Ingoglia, G., Tolosano, E. & Buehler, P. W. Haptoglobin, hemopexin, and related defense pathways—basic science, clinical perspectives, and drug development. Front. Physiol. 5, 415 (2014).

81. Pletscher-Frankild, S., Pallejà, A., Tsafou, K., Binder, J. X. & Jensen, L. J. DISEASES: Text mining and data integration of disease–gene associations. Methods 74, 83–89 (2015).

82. Grissa, D., Junge, A., Oprea, T. I. & Jensen, L. J. Diseases 2.0: a weekly updated database of disease–gene associations from text mining and data integration. Database 2022, baac019 (2022).

83. Barbour, R. et al. Red Blood Cells Are the Major Source of Alpha-Synuclein in Blood. Neurodegener. Dis. 5, 55–59 (2008).

84. Shi, M. et al. Significance and confounders of peripheral DJ-1 and alpha-synuclein in Parkinson’s disease. Neurosci. Lett. 480, 78–82 (2010).

85. Schlossmacher, M. G. et al. Detection of distinct isoform patterns of the β-amyloid precursor protein in human platelets and lymphocytes. Neurobiol. Aging 13, 421–434 (1992).

86. Feng, C. et al. Early Parkinson’s Revealed by Unlocking Longitudinal Omics at Population Scale. *medRxiv* 2026.03.12.26348299 (2026) doi:10.64898/2026.03.12.26348299.

87. Sollis, E., et al. The NHGRI-EBI GWAS Catalog: knowledgebase and deposition resource. Nucleic Acids Res. 51, D977–D985 (2022).

88. You, H. et al. Association of elevated plasma CCL5 levels with high risk for tic disorders in children. Front. Pediatr. 11, 1126839 (2023).

89. Wang, Y. et al. Micro-RNAs from Plasma-Derived Small Extracellular Vesicles as Potential Biomarkers for Tic Disorders Diagnosis. Brain Sci. 12, 829 (2022).

90. Liu, L. et al. FJX1 as a candidate diagnostic and prognostic serum biomarker for colorectal cancer. Clin. Transl. Oncol. 24, 1964–1974 (2022).

91. Shilts, J., Chen, G. & Hughey, J. J. Evidence for widespread dysregulation of circadian clock progression in human cancer. PeerJ 6, e4327 (2018).

92. Ye, Y. et al. The Genomic Landscape and Pharmacogenomic Interactions of Clock Genes in Cancer Chronotherapy. Cell Syst. 6, 314–328.e2 (2018).

93. Inoue, S. et al. Time-of-Day Immunotherapy Administration and Outcomes in Advanced Cancers. *JAMA Netw*. Open 9, e2610815 (2026).

94. Malmström, E. et al. Human proteome distribution atlas for tissue-specific plasma proteome dynamics. Cell 188, 2810–2822.e16 (2025).

95. Sun, B. B. et al. Plasma proteomic associations with genetics and health in the UK Biobank. Nature 622, 329–338 (2023).

96. Ferkingstad, E. et al. Large-scale integration of the plasma proteome with genetics and disease. Nat. Genet. 53, 1712–1721 (2021).

97. Danielson, K. M. et al. Diurnal Variations of Circulating Extracellular Vesicles Measured by Nano Flow Cytometry. PLoS ONE 11, e0144678 (2016).

98. Robles, M. S., Cox, J. & Mann, M. In-Vivo Quantitative Proteomics Reveals a Key Contribution of Post-Transcriptional Mechanisms to the Circadian Regulation of Liver Metabolism. PLoS Genet. 10, e1004047 (2014).

99. Bogaty, P. et al. Time Variability of C-Reactive Protein: Implications for Clinical Risk Stratification. PLoS ONE 8, e60759 (2013).

100. Dallmann, R., Okyar, A. & Lévi, F. Dosing-Time Makes the Poison: Circadian Regulation and Pharmacotherapy. Trends Mol. Med. 22, 430–445 (2016).

101. Dong, D., Yang, D., Lin, L., Wang, S. & Wu, B. Circadian rhythm in pharmacokinetics and its relevance to chronotherapy. Biochem. Pharmacol. 178, 114045 (2020).

102. Allan, J. S. & Czeisler, C. A. Persistence of the circadian thyrotropin rhythm under constant conditions and after light-induced shifts of circadian phase. J. Clin. Endocrinol. Metab. 79, 508–512 (1994).

103. Fan, X.-Y. et al. A meta-analysis of the value of serum TSH concentration in the diagnosis of differentiated thyroid cancer in patients with thyroid nodules. Heliyon 10, e24391 (2024).

104. Muller, J. E. et al. Circadian Variation in the Frequency of Onset of Acute Myocardial Infarction. N. Engl. J. Med. 313, 1315–1322 (1985).

105. Tofler, G. H. et al. Concurrent Morning Increase in Platelet Aggregability and the Risk of Myocardial Infarction and Sudden Cardiac Death. N. Engl. J. Med. 316, 1514–1518 (1987).

106. Cohen, M. C., Rohtla, K. M., Lavery, C. E., Muller, J. E. & Mittleman, M. A. Meta-Analysis of the Morning Excess of Acute Myocardial Infarction and Sudden Cardiac Death. Am. J. Cardiol. 79, 1512–1516 (1997).

107. Scheer, F. A. J. L. & Shea, S. A. Human circadian system causes a morning peak in prothrombotic plasminogen activator inhibitor-1 (PAI-1) independent of the sleep/wake cycle. Blood 123, 590–593 (2014).

108. Nakayasu, E. S. et al. Tutorial: best practices and considerations for mass-spectrometry-based protein biomarker discovery and validation. Nat. Protoc. 16, 3737–3760 (2021).

109. Kammers, K., Cole, R. N., Tiengwe, C. & Ruczinski, I. Detecting significant changes in protein abundance. EuPA Open Proteom. 7, 11–19 (2015).

110. Maloy, A., Alexander, S., Andreas, A., Nyunoya, T. & Chandra, D. Stain-Free total-protein normalization enhances the reproducibility of Western blot data. Anal. Biochem. 654, 114840 (2022).

111. Eriksen, H. O., Clemmensen, I., Hansen, M. S. & Ibsen, K. K. Plasma fibronectin concentration in normal subjects. Scand. J. Clin. Lab. Investig. 42, 291–295 (1982).

112. Pietzner, M. et al. Synergistic insights into human health from aptamer- and antibody-based proteomic profiling. Nat. Commun. 12, 6822 (2021).

113. Nicholas, J. C. et al. Cross-ancestry comparison of aptamer and antibody protein measures. Nat. Commun. 17, 1054 (2026).

114. Sissala, N. et al. Comparative evaluation of Olink Explore 3072 and mass spectrometry with peptide fractionation for plasma proteomics. Commun. Chem. 8, 327 (2025).

115. Niu, L. et al. Plasma proteome variation and its genetic determinants in children and adolescents. Nat. Genet. 57, 635–646 (2025).

116. Covarrubias-Pazaran, G. Genome-Assisted Prediction of Quantitative Traits Using the R Package sommer. PLoS ONE 11, e0156744 (2016).

117. Taliun, D. et al. Sequencing of 53,831 diverse genomes from the NHLBI TOPMed Program. Nature 590, 290–299 (2021).

118. Taylor-Weiner, A. et al. Scaling computational genomics to millions of individuals with GPUs. Genome Biol. 20, 228 (2019).

119. Rougemont, Q., et al. EASYstrata: an all-in-one workflow for genome annotation and genomic divergence analysis. NAR Genom. Bioinform. 7, lqaf110 (2025).

120. Purcell, S. et al. PLINK: A Tool Set for Whole-Genome Association and Population-Based Linkage Analyses. Am. J. Hum. Genet. 81, 559–575 (2007).

121. Wang, G., Sarkar, A., Carbonetto, P. & Stephens, M. A simple new approach to variable selection in regression, with application to genetic fine mapping. J. R. Stat. Soc.: Ser. B (Stat. Methodol.) 82, 1273–1300 (2020).

122. Zou, Y., Carbonetto, P., Wang, G. & Stephens, M. Fine-mapping from summary data with the “Sum of Single Effects” model. PLoS Genet. 18, e1010299 (2022).

123. Orchard, P. et al. Cross-cohort analysis of expression and splicing quantitative trait loci in TOPMed. medRxiv 2025.02.19.25322561 (2025) doi:10.1101/2025.02.19.25322561.

124. Frankish, A. et al. GENCODE: reference annotation for the human and mouse genomes in 2023. Nucleic Acids Res. 51, D942–D949 (2022).

125. Casper, J. et al. The UCSC Genome Browser database: 2026 update. Nucleic Acids Res. 54, D1331–D1335 (2025).

126. Goldfarb, T. et al. NCBI RefSeq: reference sequence standards through 25 years of curation and annotation. Nucleic Acids Res. 53, D243–D257 (2024).

127. Kuleshov, M. V. et al. Enrichr: a comprehensive gene set enrichment analysis web server 2016 update. Nucleic Acids Res. 44, W90–W97 (2016).

128. Xie, Z. et al. Gene Set Knowledge Discovery with Enrichr. Curr. Protoc. 1, e90–e90 (2021).

129. Fang, Z., Liu, X. & Peltz, G. GSEApy: a comprehensive package for performing gene set enrichment analysis in Python. Bioinformatics 39, btac757 (2022).

130. Gillespie, M. et al. The reactome pathway knowledgebase 2022. Nucleic Acids Res. 50, D687– D692 (2021).

