## Supplementary Figures for "Short-term, long-term, and genetic determinants of human plasma proteome variability"

(A)

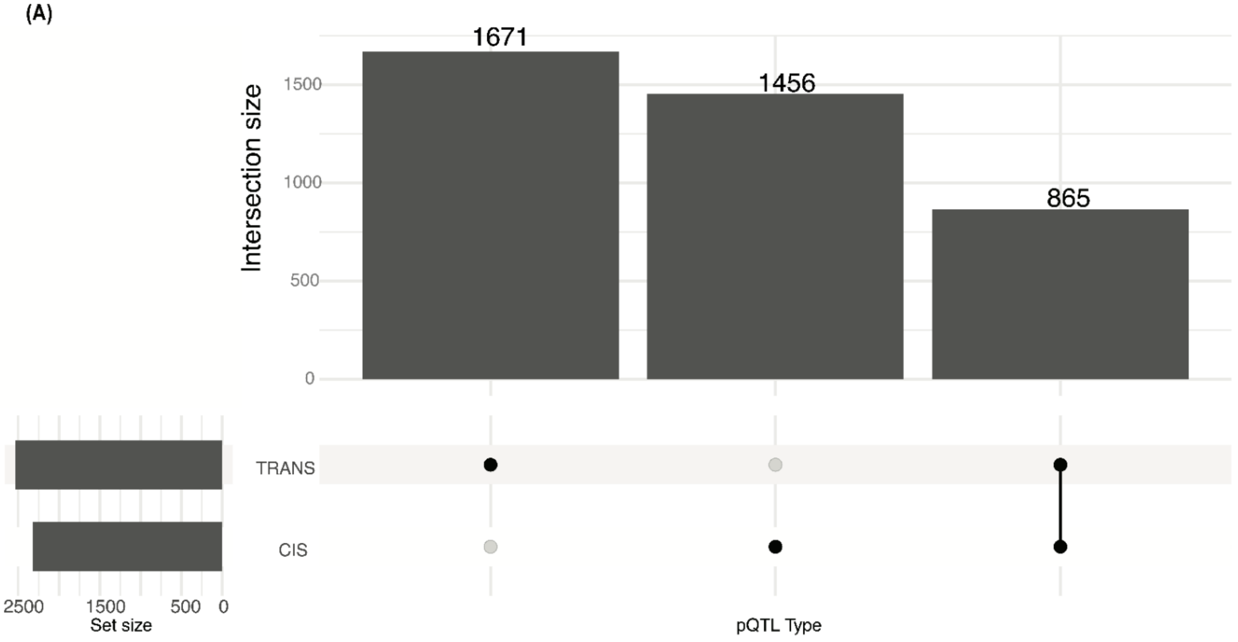


(B)

| | N SeqIDs with a Credible Set | N Credible Sets Observed | Avg N Credible Sets per SeqID | N Credible Sets with One Variant | Avg N Variants per Credible Set |
|---|---|---|---|---|---|
| *cis*-pQTL | 2,231 | 5,952 | 2.6 | 2,795 (47%) | 5.7 |
| *trans*-pQTL | 2,536 | 5,442 | 2.1 | 2,457 (45%) | 5.7 |

**Supplementary Figure 1. Summary of significant cis- and trans-pQTL credible sets observed.** (A) UpSet plot displaying the number of unique phenotypes (SeqIDs) with a significant cis-pQTL only, a significant trans-pQTL only, or a significant cis- and trans-pQTL. (B) Total counts of number of (N) SeqIDs with any significant cis- (top row) or trans- (bottom row) pQTL, total counts of cis- or trans-pQTL observed, respectively, and characteristics of the observed credible sets, including average (avg) number of credible sets per SeqID observed, number of credible sets consisting of only one variant, and average number of variants per credible set. pQTLs were calculated for SomaScan 7k protein measures in 5,149 MESA participants with linear regression under an additive model. cis-pQTL were defined as associations led by variants within 1 Megabase (Mb) of the protein-encoding gene transcription start site (TSS), considered significant at p-value<$5 \times 10^{-8}$, traditional genome-wide significance threshold. Trans-pQTL credible sets were those with variants >1Mb away from the protein-encoding gene TSS, containing at least one variant p-value<$6.8 \times 10^{-15}$ (traditional genome-wide significance threshold corrected for 7,275 SeqIDs tested).

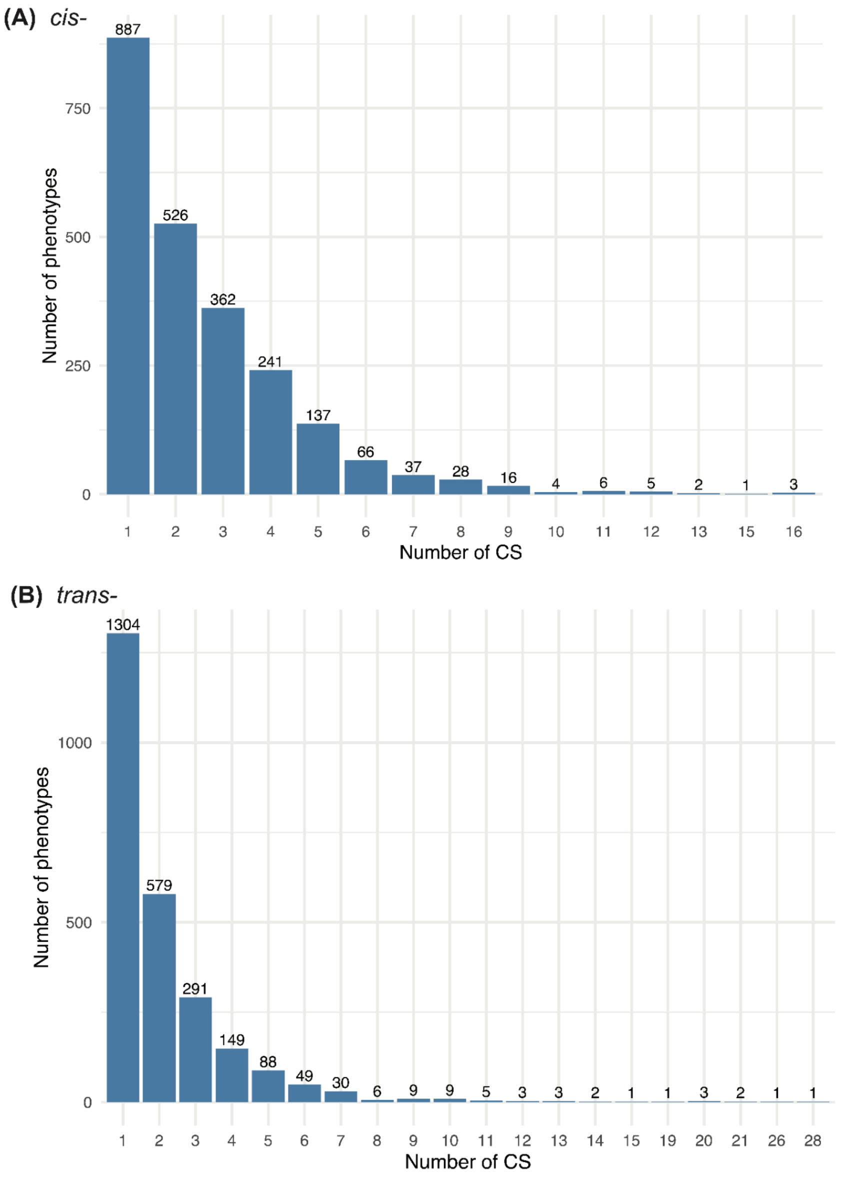


**Supplementary Figure 2. Distribution of the number of credible sets (CS) observed per phenotype (SeqID).** (A) Number of significant cis-pQTL credible sets per phenotype. Significant cis-pQTL credible sets contain at least one variant with p-value<$5x10^{-8}$ (traditional genome-wide significance threshold). (B) Number of significant trans-pQTL credible sets per phenotype. Significant trans-pQTL credible sets contain at least one variant p-value<$6.8x10^{-15}$ (traditional genome-wide significance threshold corrected for 7,275 SeqIDs tested). Note the scale of axes differs between (A) and (B).

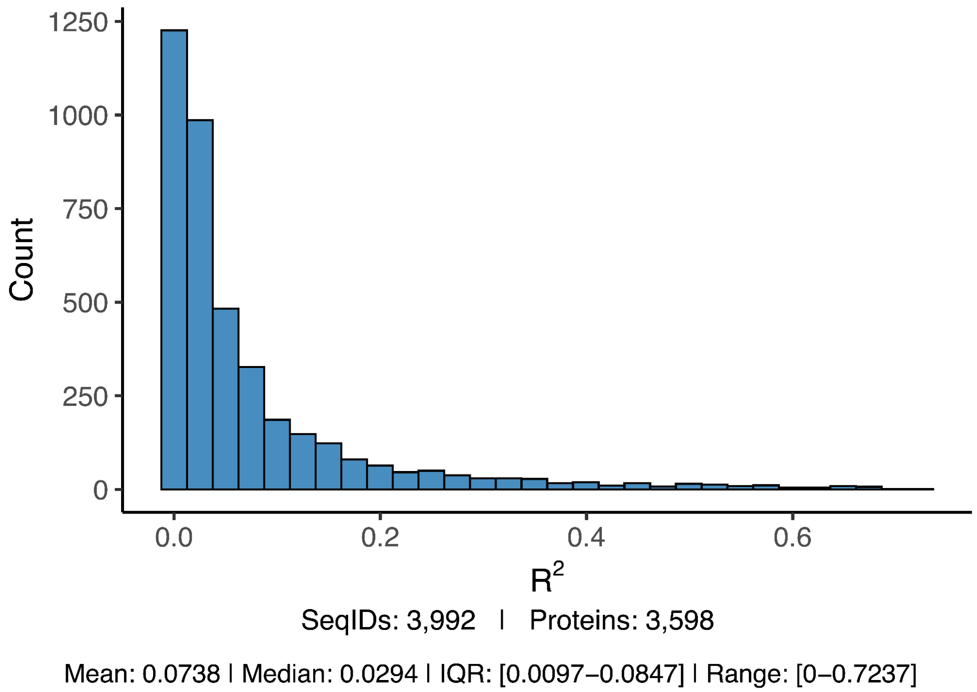


**Supplementary Figure 3. Distribution of variance explained by genetics ($R^2$), expressed as a decimal, for 3,992 SeqID measures (corresponding to 3,598 unique UniProt IDs) with at least one significant cis or trans-pQTL in 5,149 MESA participants.** Lead variants from significant cis- and trans- pQTL credible sets were used to construct a polygenic score (PGS) for each protein in MESA. Variance explained is the $R^2$ value generated from regressing protein measures against the resulting PGS. cis-pQTL were considered significant at p-value<$5x10^{-8}$, traditional genome-wide significance threshold. trans-pQTL credible sets were considered significant at p-value<$6.8x10^{-15}$ (traditional genome-wide significance threshold corrected for 7,275 SeqIDs tested). Among the 3,992 SeqIDs with a significant cis- or trans-pQTL for which GEN could be calculated, the average variance explained by pQTLs was 7.3% (median 2.9%, IQR=0.9%-8.4%).

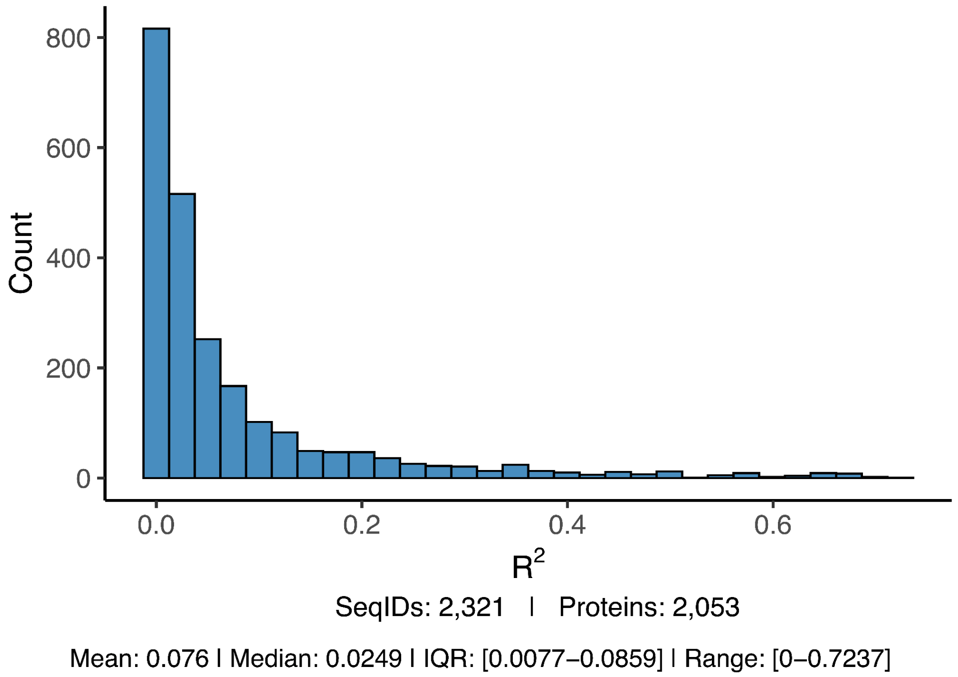


**Supplementary Figure 4. Distribution of variance explained by cis-pQTL only ($R^2$), expressed as a decimal, for 2,321 SeqID measures (corresponding to 2,053 unique proteins, as determined by UniProt IDs) with at least one significant cis-pQTL in 5,149 MESA participants.** Lead variants from significant cis-pQTL credible sets were used to construct a polygenic score (PGS) for each protein in MESA. Variance explained is the $R^2$ value generated from regressing protein measures against the resulting PGS generated using significant cis-pQTL only. cis-pQTL were considered significant at p-value<$5x10^{-8}$, traditional genome-wide significance threshold. Among the 2,321 SeqIDs with a significant cis-pQTL, the average variance explained by cis-pQTLs was 7.6% (median 2.5%, IQR=0.8%-8.6%).

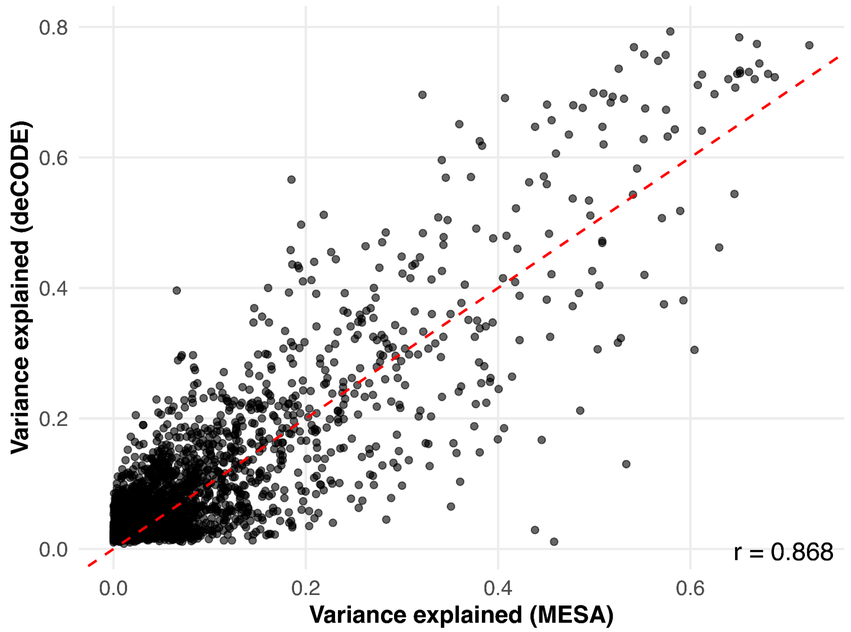


**Supplementary Figure 5. Comparison of variance explained by genetics in MESA vs. deCODE for aptamer measures with a significant pQTL in both studies.** Variance explained by genetics was compared for 2,681 aptamer measures with a significant cis- and/or trans- pQTL in 5,149 MESA participants, or with a significant cis- and/or trans- pQTL identified for SomaScan v4 measures in 35,559 deCODE Icelander participants (p-value < $1.8 \times 10^{-9}$), as reported in Ferkingstad et al (PMID: 34857953).

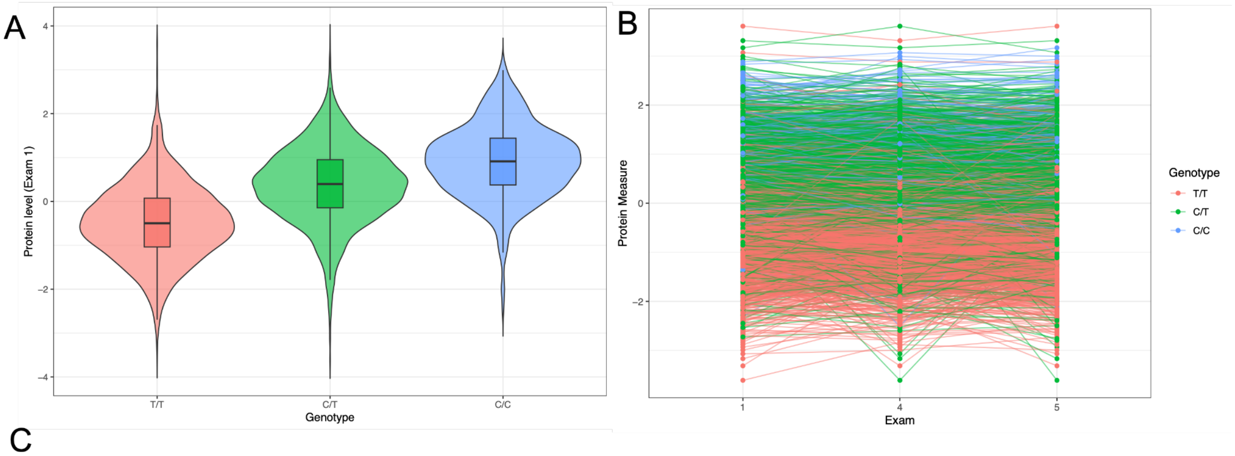


C

| SNP | Probe | Target | B | SE | P | EAF |
|---|---|---|---|---|---|---|
| chr19:54738065:T:C | seq.10428.1 | KI2S2 | 0.87 | 0.01 | $1 \times 10^{-300}$ | 0.29 |

**Supplementary Figure 6. Killer cell immunoglobulin-like receptor 2DS2 (KI2S2) *trans*-pQTL strongly associates with baseline protein level and stability across exams.** (A) Inverse normal transformed protein measure at exam 1, stratified by participant genotype at the *trans*-pQTL, in 3,274 participants with SomaScan measures at all 3 exams and whole genome sequences available. (B) Inverse normal transformed protein measures at each of the 3 MESA exams with SomaScan measures available, with lines colored by genotype of the contributing individual. (C) *trans*-pQTL association statistics for lead variant associated with killer cell immunoglobulin-like receptor 2DS2 protein measures. This variant is a non-coding variant downstream of the *KIR3DL3* gene, in strong linkage disequilibrium with a coding variant in the *KIR3DL3* gene, chr19:54738571:T:C, encoding a valine to alanine substitution. Neither the killer cell immunoglobulin-like receptor 2DS2 protein measures, nor the killer cell immunoglobulin-like receptor 3DL3 protein measures, have a *cis*-pQTL in the MESA dataset, which suggests a more complex effect than simple binding affinity.

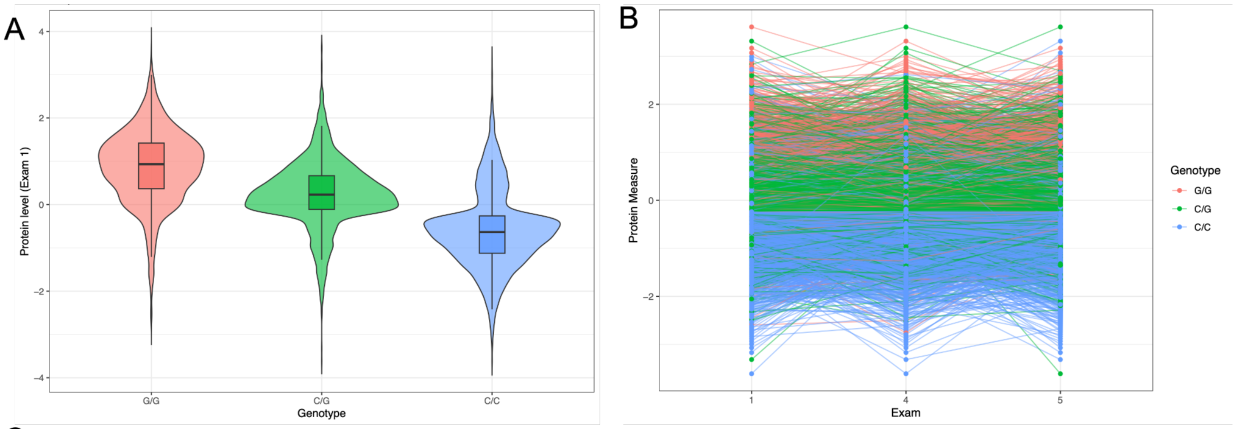


C

| SNP | Probe | Target | B | SE | P | EAF |
|---|---|---|---|---|---|---|
| chr22:30608639:G:C | seq.15560.52 | TCN2 | -0.84 | 0.02 | $1\times10^{-300}$ | 0.62 |

**Supplementary Figure 7. Transcobalamin-2 (TCN2) non-coding *cis*-pQTL strongly associates with baseline protein level and stability across exams.** (A) Inverse normal transformed protein measure at exam 1, stratified by participant genotype, in 3,274 participants with SomaScan measures at all 3 exams and whole genome sequences available. (B) Inverse normal transformed protein measures at each of the 3 MESA exams with SomaScan measures available, with lines colored by genotype of the contributing individual. (C) *cis*-pQTL association statistics for the lead variant associated with transcobalamin-2 protein measures. This is the only variant in the credible set and is an intronic variant in the *TCN2* gene.

A B

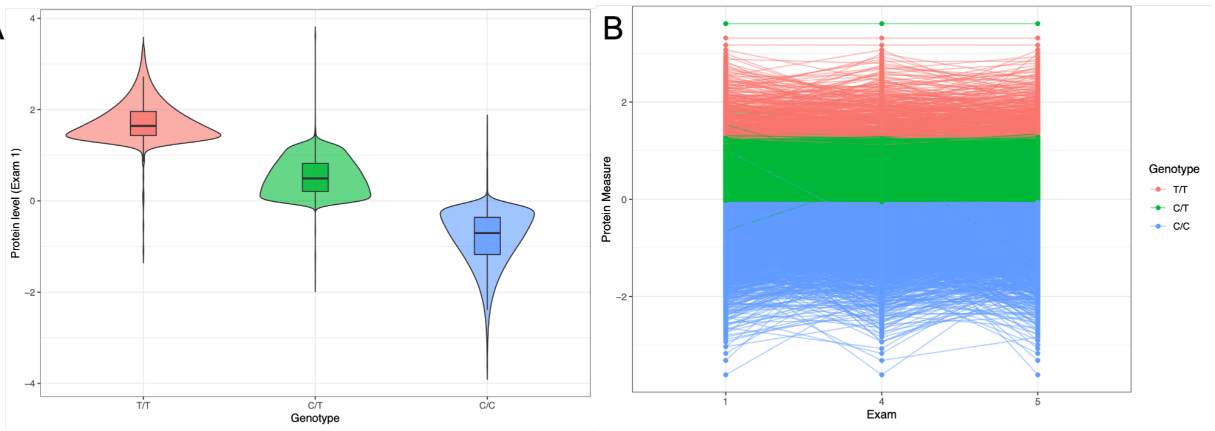


C

| SNP | Probe | Target | B | SE | P | EAF |
|---|---|---|---|---|---|---|
| chr3:186619924:T:C | seq.10966.1 | AHSG | -1.33 | 0.01 | $1x10^{-300}$ | 0.69 |

**Supplementary Figure 8. alpha-2-HS-glycoprotein (AHSG) *cis*-pQTL is driven by a missense variant that strongly associates with baseline protein level and stability across exams.** (A) Inverse normal transformed protein measure at exam 1, stratified by participant genotype, in 3,274 participants with SomaScan measures at all 3 exams and whole genome sequences available. (B) Inverse normal transformed protein measures at each of the 3 MESA exams with SomaScan measures available, with lines colored by genotype of the contributing individual. (C) *cis*-pQTL association statistics for lead variant associated with alpha-2-HS-glycoprotein protein measures. This is the only variant in the credible set and is a missense variant in the *AHSG* gene, encoding a methionine to threonine substitution.
