## Supplementary Note 1 for "Short-term, long-term, and genetic determinants of human plasma proteome variability"

The following studies, classified by system, were reviewed to identify potential biomarkers, but could not be discussed individually in the main Results section. The classifications reported here were derived from comparison with the diurnal proteomic analyses of Studies 1 and 2 described in the Methods. Of the 100 unique source-study analytes that could be mapped to targets represented in the SomaScan 7K panel, 43 (43%) showed diurnal rhythmicity, including 15 classified as circadian and 28 classified as rhythmic.

For cardiovascular diseases, Climente-González et al.[1] developed interpretable machine-learning models to predict the 10-year risk of coronary artery disease, ischemic stroke or myocardial infarction using plasma proteomic and clinical data. Among the ten most important protein predictors identified by their proteomics model, Matrix metalloproteinase-12 (MMP12) and Fas ligand (FASLG) were classified as circadian in our analysis, whereas Growth/differentiation factor 15 (GDF15) and N-terminal pro-B-type natriuretic peptide (NT-proBNP) were classified as diurnally rhythmic. In patients with stable coronary artery disease, Dregoesc et al.[2] identified six inflammatory proteins associated with the occurrence of a first major adverse cardiovascular event. Among these proteins, Fibroblast growth factor 23 (FGF23) was classified as circadian, whereas Decorin (DCN) and Tumor necrosis factor-related apoptosis-inducing ligand receptor 2 (TRAIL-R2) were classified as diurnally rhythmic. Shah et al.[3] identified 37 plasma proteins consistently associated with incident heart failure across three prospective population-based analyses and subsequently prioritized ten proteins using Mendelian-randomization analyses. Among these proteins, Neuropilin-1 (NRP1) and C-C motif chemokine 15 (CCL15) were classified as circadian, whereas Angiopoietin-like protein 3 (ANGPTL3), Follistatin-related protein 3 (FSTL3), Microfibril-associated glycoprotein 4 (MFAP4), Spondin-1 (SPON1), Sushi, von Willebrand factor type A, EGF and pentraxin domain-containing protein 1 (SVEP1) and Apolipoprotein F (APOF) were classified as diurnally rhythmic. Finally, Núñez et al.[4] identified five proteins associated with subclinical atherosclerosis and validated a three-protein panel comprising Immunoglobulin heavy constant alpha 2 (IGHA2), Apolipoprotein(a) (LPA) and Haptoglobin (HP). None of the SomaScan-mappable proteins selected from this study showed diurnal rhythmicity in our analysis.

In studies of neurological disorders and aging, Grande et al.[5] evaluated the prospective association and predictive performance of six blood biomarkers for incident all-cause and Alzheimer's disease dementia. Among these biomarkers, Neurofilament light chain (NfL) was classified as diurnally rhythmic. Hällqvist et al.[6] developed an eight-protein plasma panel to distinguish patients with Parkinson's disease and identify individuals in the premotor phase up to seven years before motor symptom onset. Among the proteins in this panel, Granulin precursor (GRN), Prostaglandin-H2 D-isomerase (PTGDS) and Dickkopf-related protein 3 (DKK3) were classified as diurnally rhythmic. Liu et al.[7] identified 13 plasma proteins associated with brain age gap and characterized nonlinear changes in the plasma proteome across brain aging. Tissue inhibitor of metalloproteinases 4 (TIMP4), Chitinase-3-like protein 1 (CHI3L1), WAP, follistatin/kazal, immunoglobulin, Kunitz and netrin domain-containing protein 1 (WFIKKN1) and Growth/differentiation factor 15 (GDF15) were classified as diurnally rhythmic. Tanaka et al.[8] identified age-associated plasma proteins predicting mortality and healthspan. Among the proteins from this analysis, Insulin-like growth factor-binding protein 4 (IGFBP4), Tissue inhibitor of metalloproteinases 1 (TIMP1), Insulin-like growth factor-binding protein 2 (IGFBP2) and Matrix metalloproteinase-12 (MMP12) were classified as circadian, whereas Stanniocalcin-1 (STC1), Cystatin-C (CST3), Cathepsin B (CTSB), Elafin/peptidase inhibitor 3 (PI3) and Growth/differentiation factor 15 (GDF15) were classified as diurnally rhythmic. Heo et al.[9] performed large-scale plasma proteomic profiling to identify proteins associated with clinical and biomarker-defined Alzheimer's disease and developed a seven-protein predictive model. Synaptic vesicle membrane protein VAT-1 homolog (VAT1) and Neuronal pentraxin receptor (NPTXR) were classified as circadian, whereas Complexin-2 (CPLX2) was classified as diurnally rhythmic.

For oncological biomarkers, Ivansson et al.[10] developed plasma-protein models to distinguish malignant ovarian tumors from benign conditions in symptomatic women. Among the proteins included in these models, WAP four-disulfide core domain protein 2 (WFDC2) was classified as diurnally rhythmic. Xing et al.[11] developed and prospectively validated a four-analyte serum panel comprising Hyaluronan-binding protein 2 (HABP2), CD163 antigen (CD163), Alpha-fetoprotein (AFP) and Protein induced by vitamin K absence or antagonist-II (PIVKA-II) for the early detection of hepatocellular carcinoma and the prediction of progression from liver cirrhosis to hepatocellular carcinoma. Hyaluronan-binding protein 2 (HABP2) and PIVKA-II were classified as diurnally rhythmic. However, PIVKA-II is an undercarboxylated form of prothrombin, and its mapping to a prothrombin/F2 SomaScan target does not demonstrate PIVKA-II-specific diurnal rhythmicity. Finally, Liu et al.[12] investigated Four-jointed box kinase 1 (FJX1) as a candidate serum biomarker for the diagnosis and prognosis of colorectal cancer. FJX1 was classified as circadian.

In inflammatory, infectious, pulmonary and psychosocial contexts, He et al.[13] characterized plasma-protein changes preceding rheumatoid arthritis and developed protein-based models predicting responses to methotrexate combined with either leflunomide or hydroxychloroquine. Among the seven proteins used in these treatment-response models, Carbonyl reductase 1 (CBR1) and Collagen alpha-1(I) chain (COL1A1), both included in the methotrexate-leflunomide model, were classified as circadian. Shu et al.[14] developed an 11-protein biomarker set and multivariable models to distinguish COVID-19 severity and predict clinical outcomes. None of the SomaScan-mappable proteins in this biomarker set showed diurnal rhythmicity in our analysis. The discovery cohort included serial samples collected at different stages of disease, with up to four samples from each fatal case and two samples from each survivor with mild or severe disease. Although the clock time of collection was not reported, if these clinically collected samples were distributed across the day, repeated sampling may have averaged part of the diurnal variation across the dataset and reduced the probability that strongly diurnally rhythmic proteins were selected by the model. This provides a plausible explanation for the absence of overlap, but does not demonstrate that the study formally controlled for sampling time. Shen et al.[15] characterized plasma-proteomic signatures of social isolation and loneliness and identified five proteins with Mendelian-randomization evidence of an association with loneliness. Among these proteins, Fatty acid-binding protein 4 (FABP4) was classified as circadian, whereas Asialoglycoprotein receptor 1 (ASGR1) and GDNF family receptor alpha-1 (GFRA1) were classified as diurnally rhythmic. Finally, Suryadevara et al.[16] integrated transcriptomic, alternative-splicing and plasma-proteomic data to identify biomarkers of computed-tomography-defined emphysema and develop predictive biomarker models. PC3-secreted microprotein (PSMP; encoded by MSMP) was classified as circadian, whereas Receptor-type tyrosine-protein phosphatase delta (PTPRD) was classified as diurnally rhythmic.

Together, these observations indicate that diurnal rhythmicity occurs among biomarkers selected for diverse clinical and biological outcomes and is not restricted to a specific disease area or proteomic platform.

1. Climente-González, H. *et al.* Interpretable machine learning leverages proteomics to improve cardiovascular disease risk prediction and biomarker identification. *Commun. Med.* **5**, 170 (2025).
2. Dregoesc, M. I. *et al.* Relation Between Plasma Proteomics Analysis and Major Adverse Cardiovascular Events in Patients With Stable Coronary Artery Disease. *Front. Cardiovasc. Med.* **9**, 731325 (2022).
3. Shah, A. M. *et al.* Large scale plasma proteomics identifies novel proteins and protein networks associated with heart failure development. *Nat. Commun.* **15**, 528 (2024).
4. Núñez, E. *et al.* Unbiased plasma proteomics discovery of biomarkers for improved detection of subclinical atherosclerosis. *eBioMedicine* **76**, 103874 (2022).
5. Grande, G. *et al.* Blood-based biomarkers of Alzheimer's disease and incident dementia in the community. *Nat. Med.* **31**, 2027–2035 (2025).
6. Hällqvist, J. *et al.* Plasma proteomics identify biomarkers predicting Parkinson's disease up to 7 years before symptom onset. *Nat. Commun.* **15**, 4759 (2024).
7. Liu, W.-S. *et al.* Plasma proteomics identify biomarkers and undulating changes of brain aging. *Nat. Aging* **5**, 99–112 (2025).
8. Tanaka, T. *et al.* Plasma proteomic biomarker signature of age predicts health and life span. *eLife* **9**, e61073 (2020).
9. Heo, G. *et al.* Large-scale plasma proteomic profiling unveils diagnostic biomarkers and pathways for Alzheimer's disease. *Nat. Aging* **5**, 1114–1131 (2025).
10. Ivansson, E. *et al.* Large-scale proteomics reveals precise biomarkers for detection of ovarian cancer in symptomatic women. *Sci. Rep.* **14**, 17288 (2024).
11. Xing, X. *et al.* Proteomics-driven noninvasive screening of circulating serum protein panels for the

early diagnosis of hepatocellular carcinoma. *Nat. Commun.* **14**, 8392 (2023).
12. Liu, L. *et al.* FJX1 as a candidate diagnostic and prognostic serum biomarker for colorectal cancer. *Clin. Transl. Oncol.* **24**, 1964–1974 (2022).
13. He, S. *et al.* A longitudinal cohort study uncovers plasma protein biomarkers predating clinical onset and treatment response of rheumatoid arthritis. *Nat. Commun.* **16**, 6692 (2025).
14. Shu, T. *et al.* Plasma Proteomics Identify Biomarkers and Pathogenesis of COVID-19. *Immunity* **53**, 1108-1122.e5 (2020).
15. Shen, C. *et al.* Plasma proteomic signatures of social isolation and loneliness associated with morbidity and mortality. *Nat. Hum. Behav.* **9**, 569–583 (2025).
16. Suryadevara, R. *et al.* Blood-based Transcriptomic and Proteomic Biomarkers of Emphysema. *Am. J. Respir. Crit. Care Med.* **209**, 273–287 (2024).
